# The population dynamics of a canonical cognitive circuit

**DOI:** 10.1101/516021

**Authors:** Rishidev Chaudhuri, Berk Gerçek, Biraj Pandey, Adrien Peyrache, Ila Fiete

## Abstract

The brain constructs distributed representations of key low-dimensional variables. These variables may be external stimuli or internal constructs of quantities relevant for survival, such as a sense of one’s location in the world. We consider that the high-dimensional population-level activity vectors are the fundamental representational currency of a neural circuit, and these vectors trace out a low-dimensional manifold whose dimension and topology matches those of the represented variable. This manifold perspective — applied to the mammalian head direction circuit across rich waking behaviors and sleep — enables powerful inferences about circuit representation and mechanism, including: Direct visualization and blind discovery that the network represents a one-dimensional circular variable across waking and REM sleep; fully unsupervised decoding of the coded variable; stability and attractor dynamics in the representation; the discovery of new dynamical trajectories during sleep; the limiting role of external rather than internal noise in the fidelity of memory states; and the conclusion that the circuit is set up to integrate velocity inputs according to classical continuous attractor models.

---

It has long been clear that the brain represents sensory, motor, and internal variables in distributed codes across large populations of neurons. In turn, theoretical models of computation in the brain have emphasized that neural circuit dynamics must be understood in terms of the emergence of simple structures from the collective interactions of large numbers of neurons^1–8^, and that robust representation and memory involve the formation of low-dimensional attractors in the population dynamics.

Until relatively recently, experimental techniques permitted access to only one or a few neurons at a time, but simultaneous recordings of multiple neurons are making feasible the theoretically suggested approach of characterizing the structure and dynamics of neural responses at the population level. This approach is beautifully illustrated in recent demonstrations of low-dimensional trajectories in sensory and motor circuits^9–12^.

Our work proceeds from four central premises: 1) In distributed codes, information representation, computation, and dynamics unfold at the level of the neural population and the collective states of a circuit are the natural way to understand them. 2) If a circuit represents a low-dimensional variable of given dimension and topology, the high-dimensional states of the circuit will be localized to a low-dimensional subspace or manifold of matching dimension and topology. 3) Characterizing the structure of this manifold can enable unsupervised discovery and decoding of the internally coded (latent) variable. 4) Examining manifold structure and dynamics on and off the manifold across a range of behavioral states as the inputs to the circuit change can reveal aspects of circuit mechanism.

We illustrate a method to characterize the manifold structure of data, use this characterization to discover, in a blind or unsupervised way, low-dimensional internal states, provide a blind time-resolved decoding of these states, and provide support for the predictions of a classic mechanistic circuit model, using the mammalian head direction system as our subject. The head direction (HD) system in mammals and insects^13–21^ is a high-level cognitive circuit that uses various external and internal cues to compute an estimate of the direction of heading of the animal with respect to the external world. It is a proving-ground for the manifold-based approach to unsupervised discovery of the encoded variables because it represents internal cognitive states that need not directly reflect externally measured variables during waking, and this dissociation between internal and external states holds even more true during sleep (as we will see). At the same time, the HD system also helps us illustrate how a manifold approach can yield genuinely new insight into the structure, dynamics, and mechanisms of a long-studied neural circuit.

Two decades ago, theoretical models^4;22–25^ of the HD circuit predicted the existence of a stable, one-dimensional (henceforth 1D) ring-shaped stable manifold in the high-dimensional state space of the population activity, a more abstract and fundamental feature than details about the shapes of tuning curves, connectivity profiles, or physical placements of neurons in the circuit. The notion of stability means that perturbations of any kind in the high-dimensional space away from the ring should quickly and preferentially flow back to the ring. If the HD circuit is an integrator, then the states along the ring and changes in state along the ring for equivalent changes in the represented variable should in some sense be equal. The basic elements of the HD circuit models have since been extended to explain the dynamics of other neurons, including grid cells^7^. The same circuit models can further explain how, through the integration of a different velocity signal, the brain could form representations in more abstract metric spaces^26;27^. Thus, testing the extent to which these models are really correct descriptors of circuit mechanism is a question of broad importance.

So far, pairwise correlations between mammalian HD cells^15;28–32^ and a topographically ordered *physical* HD-coding ring structure in flies^18–21^ are consistent with the hypothesized models. Here we show directly that the long-hypothesized low-dimensional state-space ring structure and attractive dynamics can be directly visualized in the population responses of the mammalian HD system. The dynamics revealed in this circuit during sleep states provide new evidence about the mechanisms that allow these states to be maintained and updated.

Portions of these results have been presented at conferences^33;34^.

## Results

The instantaneous (temporally binned) response of *N* neurons is a point in an *N*-dimensional *state space* where each axis represents the activity of one neuron (Fig. 1a). The collection of such snapshots of the population activity forms a cloud in the state space. If the population encodes some variable of dimension *D_m_* ≪ *N* and a certain topology, the point cloud should trace out a manifold of the same dimension and topology, although the shape may be convoluted. In what follows, we describe how characterizing the manifold’s topology and structure, then analyzing dynamics on the manifold, can permit us to extract the latent encoded variables in an unsupervised way and deduce key aspects of circuit mechanism.

**Figure 1:**
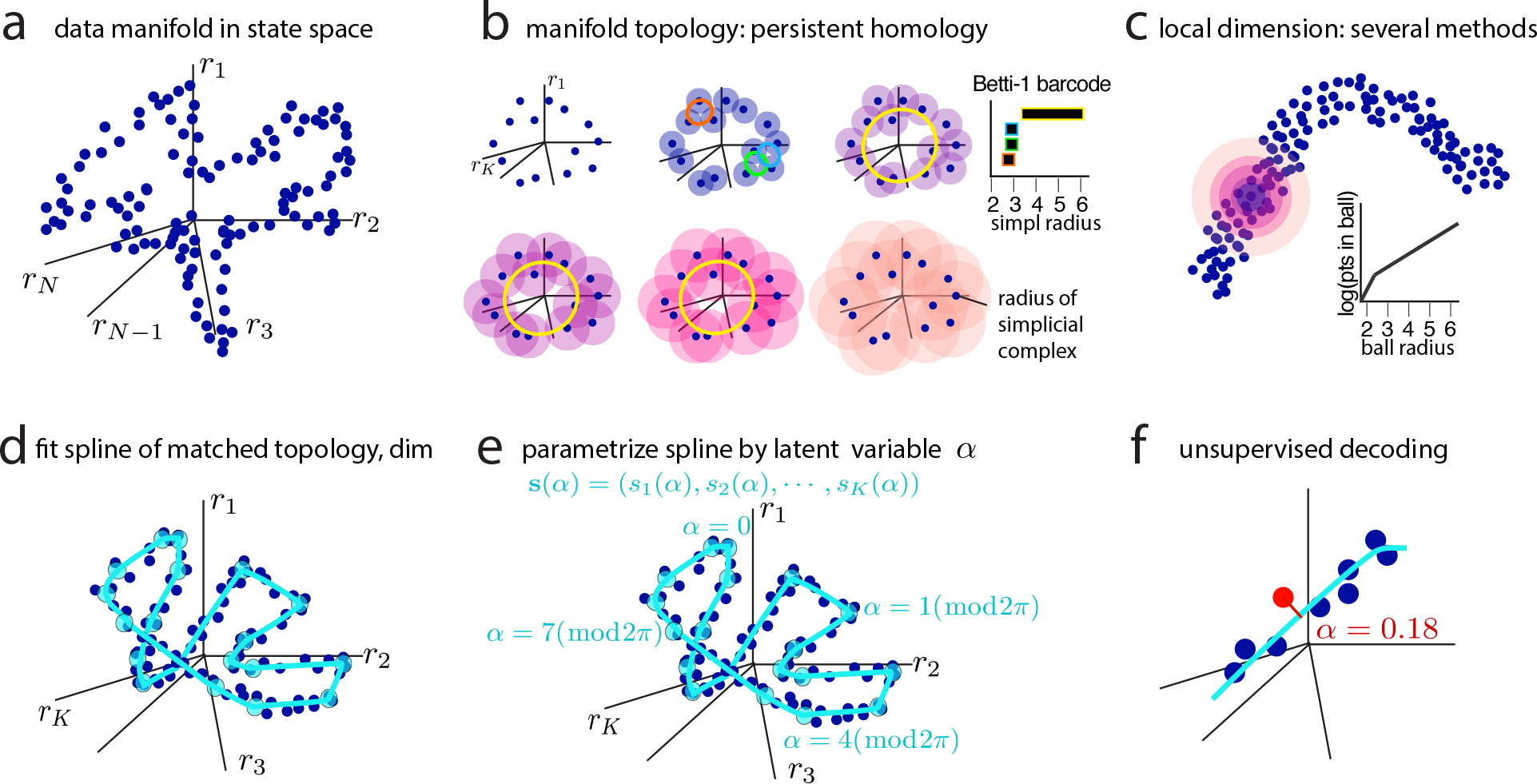
Population activity as a manifold and a method for manifold characterization. (a) Each blue point is a vector representing instantaneous activity in the neural population: vector component (*r_i_*) is the activity of the *i*th neuron. The points form a manifold. (b) Persistent homology to determine topology for general manifolds: Balls of different size centered on the datapoints represent different simplicial radii (*r*) or scales. At a given radius, sets of connected points form simplicial complices (SI S2 for details). Each complex is characterized by a set of Betti numbers; depicted here are the loop features of the complices (colored rings), at the different scales. Loops appear in panel 2 and disappear (are filled in) by panel 3; another appears in panel 3 and persists until the last; it is deemed significant because of its persistence. Inset: Betti-1 barcode, or the range of radii over which each loop persists. (c) Local dimension can be estimated by several methods. Schematic shows correlation dimension: Count the number of points in a ball of radius *r* centered at a point on the manifold, as a function of *r*. The number of points should grow as 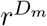 if the manifold is *D_m_* dimensional. (d) The data manifold is fit by a spline (cyan line) of dimension and topology as determined in (a-c). The spline uses a few anchor points (cyan circles) determined by clustering methods, with connecting polynomial curves. (e) The spline is parametrized by assigning coordinates along its length. The coordinates represent the values of an internal (latent) state that the circuit is assumed to encode. (f) Moment-by-moment decoding of the internal state is done by reading out the parametrization value at the point on the spline closest to the datapoint.

### Spline parameterization for unsupervised decoding (SPUD)

We characterize the global topology of the manifold using methods of topological data analysis, specifically persistent homology^35^: The method starts from the point-cloud of data, Fig. 1a, blurring the points at different resolutions or scales, and at each resolution examining the emergence of connected groups of data points called simplicial complexes, Fig. 1b. A simplicial complex can contain certain structures within it, such as a ring or a torus and so on. Betti numbers form a list of binary structural designations that characterize the complexes: A complex that is (topologically) equivalent to a flat, holeless sheet of some dimension has only a non-zero Betti-0 number; all higher-order Betti numbers are zero. A (convoluted) ring or a cylinder, each of which enclose a single hole, have a non-zero Betti-0 number and Betti-1 number, but none of higher-orders; a figure-8 shaped complex will have a non-zero Betti-0 number and two Betti-1 numbers, but none of higher order; a (hollow) torus, which contains two circular holes and one two-dimensional void, has a non-zero Betti-0 number, two Betti-1 numbers and one Betti-2 number; and so on for more complex objects. In noisy data, if a Betti number for a structure persists over many scales (Fig. 1b), this feature is robust against noise and thus deemed significant. Topological data analysis prescribes how to characterize the Betti numbers of complexes in a dataset^35^.

To decode the internal state encoded by the manifold, we perform the following steps (details in Methods): 1) Consider the binned spiking data as points in a high-dimensional state space, Fig. 1a; 2) Determine the topology of the point cloud using methods of topological data analysis (TDA), specifically persistent homology^35^, Fig. 1b, and the introduction of a neighborhood-thresholded TDA (nt-TDA) for increased robustness to noisy data (see SI S2.2); 3) estimate intrinsic local manifold dimension using various methods including correlation dimension^36^, Fig. 1c; 4) Fit the manifold with a spline of matching topology and dimension, Fig. 1d; 5) Parametrize the spline by a smoothly changing variable of matching dimension and topology, Fig. 1e; the resulting parametrization is interpreted as the values of the encoded latent variable or internal state. 6) Given a population state at any moment in time, decode that state by projecting it to the nearest point on the spline; the parametrization value at that point is the unsupervised estimate of the value of the encoded latent variable, Fig. 1f. In this entire decoding process, when the manifold is topologically non-trivial in structure, constructing a low-dimensional embedding is neither *necessary* (though nonlinearly embedding the manifold into some intermediate-dimensional space can be practically useful for efficient spline fitting, for modestly improving the spline fits because the embedded manifold has fewer narrow kinks or convolutions (SI Fig. S4 to see the modest gains from dimensionality reduction when one is data-limited), and for visualization when the manifold dimension is sufficiently small), nor is it *sufficient*. Dimensionality reduction provides a global coordinate system that is not sufficient for obtaining a minimal parameterization for manifolds that are not topologically equivalent to some hole-free *K*-dimensional sheet: For instance, the minimum embedding dimension for a 1D ring is 2D, and dimensionality reduction will yield at best a global 2D parametrization of the 1D variable, which represents a failure to discover the real 1D latent variable. This problem is general for all topologically non-trivial manifolds, because there is an important difference between global and local coordinates, and global coordinates provide either a non-unique or too-high dimensional parametrization of the manifold. The on-manifold parameterization method we describe yields a local rather than global coordinate system to describe the manifold, and extracts the correct one-dimensional structure of the encoded variable.

### Ring manifold and unsupervised decoding in the mammalian HD system

We apply SPUD to neural activity data recorded from the anterodorsal thalamic nucleus (ADn) of mice that are awake and foraging in an open environment along random, variable paths with variable velocities, as well as during intervening REM and nREM periods^32^. Note that the animal’s waking behavior is not low-dimensional: Unlike in many other applications of manifold methods^10;12^, the animal is not constrained to move along specific trajectories or perform a stereotyped task. Further, during sleep the neural dynamics are not known to be externally constrained in any way.

We show manifolds and compute Betti barcodes for the best session in each of the 7 recorded animals (Figs. S1, S7 and S11). All remaining results are based on the (3/7) animals where the RMS difference between measured and decoded angle during waking is < 0.5 rad. For any session that we use for unsupervised decoding, we include all recorded thalamic cells, with no sub-selection based on tuning. For each cell, instantaneous firing rates are obtained by convolution with a Gaussian kernel (100ms standard deviation). The first problem is to determine whether the data exhibit some simple low-dimensional manifold structure in state space. The states of the network during waking exploration appear — through direct visualization of a nonlinear low-dimensional embedding from the high-dimensional state space^37^^a^ - to lie on a strikingly low-dimensional albeit highly non-linear manifold, in the form of a convoluted ring, Fig. 2a (see Fig. S1 for all 7 animals in the data set, and Supplementary Movie 1 for a 3D view). Because waking behavior is not low-dimensional, the low-dimensional structure we see is intrinsic, rather than imposed by the environment or task.

**Figure 2:**
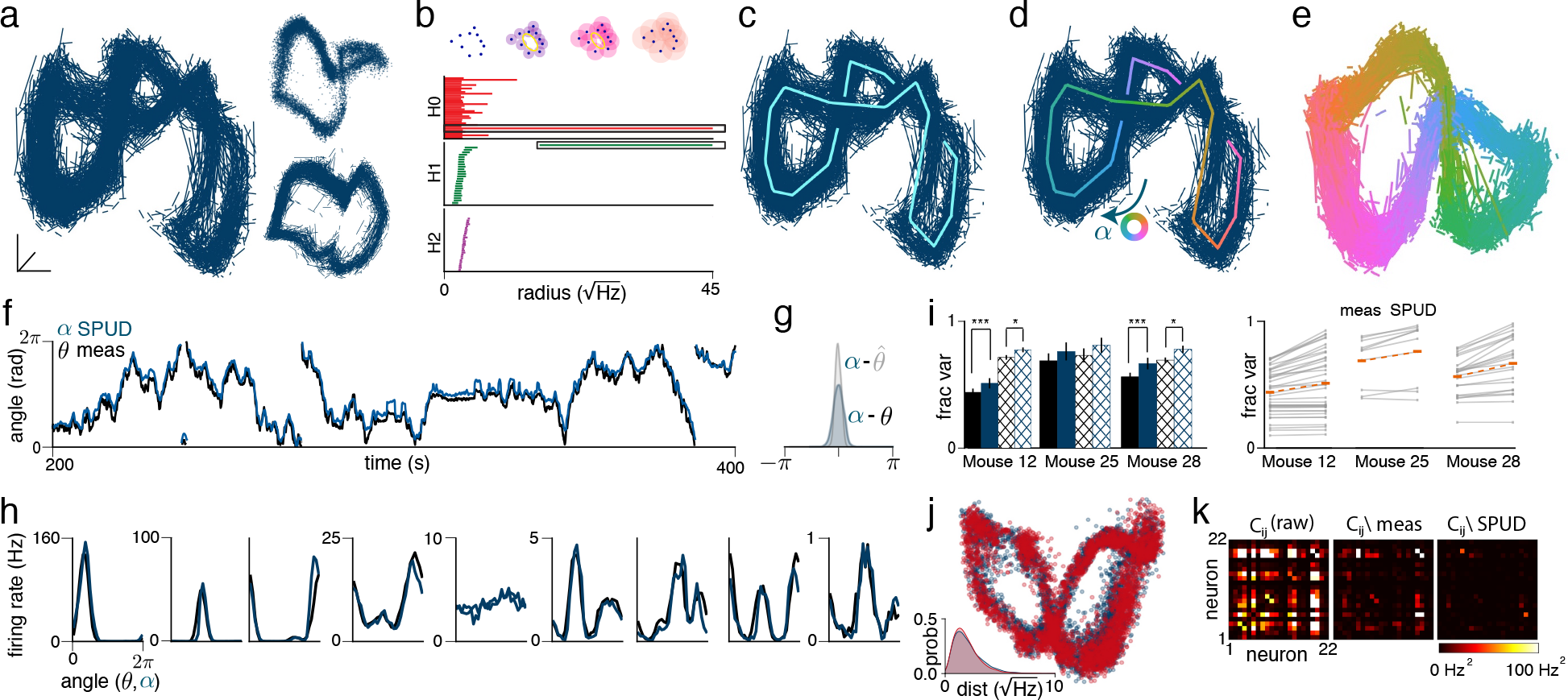
Unsupervised discovery and time-resolved decoding of encoded variable through manifold characterization. Throughout, *θ* = direct measurement of the animal’s head orientation from LED tracking; 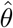 = supervised decoder’s estimate of the brain’s representation (using tuning curves); *α* = unsupervised latent variable estimate. This figure: ADn data from single animal for full waking episode (31 minute interval). (a) Visualization of manifold (by Isomap^37^), with every alternate pair of temporally adjacent points connected. Inset shows point cloud (top) and alternate view of manifold (bottom). Note that PCA often fails to pull out the ring structure of manifold, Fig. S2. (b) Betti-0, −1 and −2 barcodes: Simplicial complex radii on abscissa (schematic at top: complexes constructed from data at different radii): the start and end of a horizontal bar in the middle plot signals the appearance and disappearance of some ring (a non-zero Betti-1 feature) in the data at the corresponding radii. Long bar: a ring that appears at 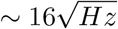 and persists until 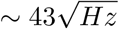 (for description of analysis and units see S2). (c) Spline fit to point cloud. (d) Parametrization of spline by coordinate *α* (arbitrary origin). (e) Coloring of neural states by unsupervised latent variable estimate (i.e., *α*). (f) Comparison of *α* and *θ*. (The origin and direction around the ring for measured head angle and for unsupervised decoding, both arbitrary choices, are matched to facilitate comparison, only after unsupervised decoding is complete.) (g) Histogram of differences between *α* and *θ* and between *α* and 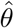. (h) Fully unsupervised tuning curve estimate (blue) versus supervised tuning curve estimate (black). (i) Left: Average fraction of variance explained by *θ* (black) and *α* (blue) under a Poisson spiking model (solid bars) and an overdispersed (hatched bars) model. Error bars show standard error; significance is from two-sided binomial test (see SI S5). Right: Variance explained for individual cells under Poisson model, with means shown in orange. (j) Manifold from data (blue) and from an overdispersed spiking model (red), with overdispersion estimated from the data (see SI S5) and applied as uncorrelated across neurons. Inset shows distribution of distances from manifold fit for data and model. (k) Covariance of firing rates (left) and covariance conditioned on either *θ* or *α*.

To independently verify that the structure visualized in the embeddings is real rather than an artefact of the visualization process, and that important, higher-dimensional and topological structures are not lost, we turn to topological data analysis, in particular the persistent homology of simplicial complexes^35^. Topological methods are more general because they permit characterization of manifolds that are topologically non-trivial in structure and higher-dimensional^35^, when direct visualization is not possible.

From the persistent homology of simplicial complexes constructed from the waking data (see Methods), we confirm an open loop or ring structure in the data that persists over several spatial scales (H1 plot, Fig. 2b), and also find no evidence of a toroidal or more complex topological structure (H2 plot, Fig. 2b; contrast Figure S3). As shown below, there is no evidence of additional structure down to the resolution of the data.

With the confirmation of a ring topology, we fit to the manifold a nonlinear spline with the same topology (Fig. 2c) and then isometrically parameterize the spline along its length with a circular variable whose values are indicated by the color of the spline (Fig. 2d). Points on the manifold are colored according to the nearest parameterization value (Fig. 2e).

Strikingly, the decoded latent variable very closely matches (up to an arbitrary choice of origin and direction) the directly measured head angle (from LED’s on the animal’s head), Fig. 2f,g (see Fig. S4 for other animals). The match serves as a direct validation that the extracted ring structure is real, and of the hypothesis that the topology of neural representations should match the topology of the represented variables.

It was not clear *a priori* that an isometric parameterization would suffice for this level of accuracy in decoding. The fact that isometric parameterization along the neural population response manifold produces excellent decoding implies that equal amounts of neural code length or activity variation are given to equal changes in head angle and that therefore no specific angles are favored for greater representational resources than others, exactly consistent with the expectations for an head velocity integrator circuit, in which all states have to be equivalently represented and equivalently changeable so that a unit velocity input produces a unit change in represented angle regardless of the angle. This *isometry property* enables accurate integration of a velocity signal regardless of starting angle.

Next, we regress the time-varying firing rates of individual cells onto the latent variable estimate. This allows us to recover neural tuning curves in a fully blind way, Fig. 2h. The unsupervised tuning curves capture 71% ± 2.8% of the variance of tuning curves constructed the traditional, supervised way (Fig. S5). Relatively flat tuning curves are also consistent across the supervised and unsupervised estimates under the assumption of an encoded variable that is one-dimensional and circular; cells with flatter curves are slightly but not significantly more overdispersed in their spiking relative to well-tuned cells (correlation between variance of tuning curve and overdispersion is 0.25, *p* of 0.12; data not shown).

The unsupervised latent variable estimate appears to more faithfully track the internal representation of head angle than it does the measured HD, assessed from the even closer match between the latent variable estimate and internal state estimate constructed from a supervised (tuning-curve-based) decoder of HD circuit activity (Fig. 2g; Fig. S4, S5 for other animals). Indeed, the measured HD is not guaranteed to accurately report the animal’s internal representation, which may slip relative to the measured HD for various reasons, including the possibility that the animal is uncertain about its HD, is representing past or future HD states, or because of errors in the experimental measurement of HD. Further, our latent variable estimate explains more of the variance of neural spiking (cross-validated, Fig. 2i) than does the measured HD, a confirmation that the unsupervised latent variable estimate is a more accurate reflection of the internal representation of HD (see Fig. S5 for further controls including leave-one-cell-out analyses where the variance of a neuron is explained using SPUD and activity in the rest of the population).

A natural question is whether the waking manifold encodes additional variables not yet discovered, for example in the thickness of the ring. With a finite signal-to-noise ratio (SNR) in the dataset it is impossible, with any method, to exclude the possibility of structure in the data that is significantly smaller than the noise. However, we may search for additional coding structure down to the resolution limit imposed by noise, by asking whether the data exhibits a spread or structure not explained by the 1D ring structure together with independent neural spiking noise. We thus generate synthetic data based only on the 1D latent variable extracted through SPUD, with spikes generated independently (after conditioning on the 1D variable) using an independent point process per cell with dispersion matched empirically to the data. The structure and spread of the resulting point cloud closely match those in the real data, Fig. 2j, suggesting that the circuit is not encoding additional variables in the form of shared (correlated) structure in other dimensions of its response. (By contrast, we do find an additional coding dimension in postsubicular HD cells (coding for head velocity; data not shown) as well as in ADn during nREM sleep (shown later)).

We search further for small-scale cooperative coding beyond the 1D ring in the population response by directly examining patterns of covariation between neurons after removing the effects of their shared angular coding around the ring manifold, Fig. 2k. Patterns of correlation are strong before conditioning on the angular variable, but weak after (the ratio of the Frobenius norm of the residual covariance matrix to the norm of the raw covariance matrix is only 6% when conditioning on the unsupervised latent variable estimate, suggesting that it captures ~ 94% of the data covariance; by contrast, the ratio after conditioning on measured HD is 25%, consistent with earlier findings that measured HD is a worse indicator of internal state than our unsupervised estimate). These results demonstrate that there is little discernible additional structure in the waking manifold, and the ADn appears to support the encoding of only a single one-dimensional circular variable during waking, down to the resolution (SNR) of the present data.

The SNR in these data and results is primarily limited by the number of simultaneously recorded cells. Larger samples of simultaneously recorded neurons will improve SNR and reduce the scatter around the ring, allowing discovery of finer additional structure or further downgrading the possibility that it exists.

### The ring manifold is autonomously generated and attractive

Above, we highlighted how regarding neural population responses as lying on a manifold and then characterizing the structure of the manifold can permit unsupervised decoding of latent variables. In what follows, we show how the manifold view reveals the collective dynamics of the circuit, in a direct, easy, and natural way. First, we will consider coarse-grained net flows of states on and off the manifold. Second, we will consider the fine time-resolved dynamics of trajectories on the manifold.

Through these analyses, we test the key predictions^1;3;4;7^ of continuous attractor network models (properties 1-4) and models of neural integrators for continuous variables (properties 1-5), Fig. 3a (see Fig. S6 for a network model): 1) The high-dimensional network response occupies a low-dimensional continuum of states with dimension and topology matching those of the encoded variable(s). 2) The states are autonomously generated and stabilized, and capable of self-sustained activation when sensory inputs are removed. 3) The manifold is an attractor: states initialized away from the manifold flow rapidly back toward it. 4) The states along the manifold are energetically equal, with no flux or net flow along the manifold. 5) A velocity input, encoding the time-derivative of the represented variable, drives the circuit in a special direction, specifically, along the low-dimensional manifold in the high-dimensional state space. Note that these predictions are fundamentally in terms of the population manifold, hence most naturally tested at that level, when the data make it possible.

**Figure 3:**
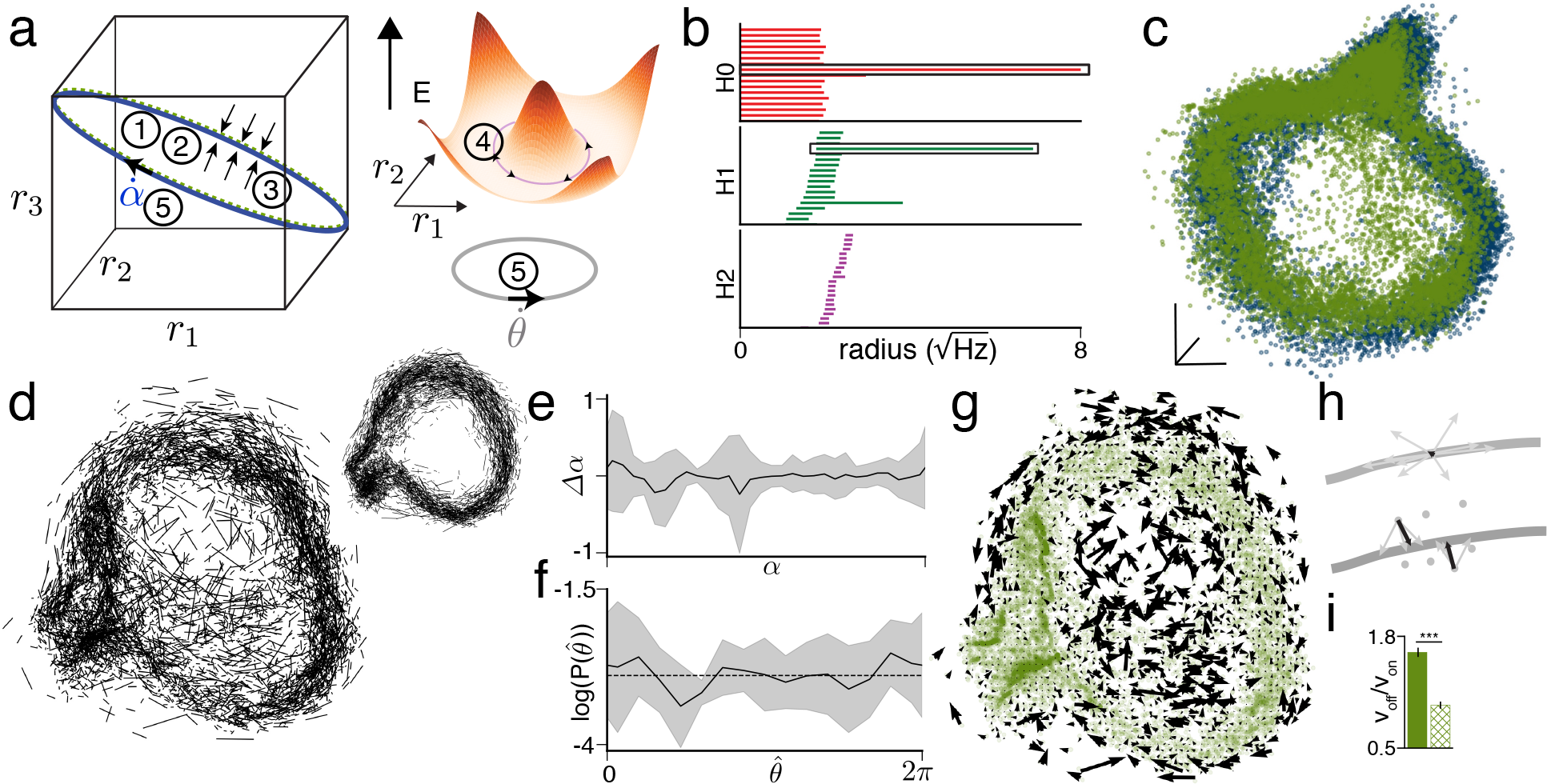
REM sleep states, fluxes, and dynamics suggest that the manifold is internally generated and attractive. (a) Schematic of attractor model predictions (see main text for list). (b) Betti-1 barcode reveals 1D ring structure preserved during REM sleep. (c) Joint visualization of REM (green) and waking data (dark blue) using Isomap^37^. (d) Flows along REM manifold: each bar represents a velocity vector at a moment in time. Inset shows wake manifold. (e) Single session mean and standard deviation of change in decoded angle as a function of angle. (f) Mean and standard deviation of angle occupancy (from tuning curve decoder) across sessions. (g) Flux on and off manifold. (h) Schematic showing that flux should be small on manifold, because velocity vectors tend to point in both directions along manifold and thus average out (top panel), while flux off manifold should be large, because velocity vectors tend to point towards manifold (bottom panel). (i) Ratio of size of flux vector off manifold to flux vector on manifold. Error bars show standard deviation; hatched bar shows shuffled control (*p* < 10^−6^; see SI S8).

Results above already directly support property 1). To study autonomous dynamics, we examine the states of the circuit during sleep, when the circuit no longer receives spatial or directional input from the world. During REM sleep, the states again lie on a ring, Fig. 3b (and Supplementary Movie 2), and moreover are essentially identical to those from awake exploration, laying on the same ring manifold, Fig. 3c (Fig. S7 for other animals). The result is in direct support of property 2), and consistent with a similar previous conclusion inferred from preserved pairwise correlations during sleep^32^.

### States on the manifold are energetically flat or equivalent

To test the equivalence of states along the manifold, we examine the coarse occupancy and dynamics of manifold states during REM sleep, when the circuit is not driven by the external world (spatial exploration could be biased). First, we construct instantaneous (undirected) vectors or bars linking states at adjacent time-points, Fig. 3d; the length of a bar is proportional to the mean speed at that time. If the manifold contained a few prominent sinks or discrete fixed points, there would be fast flow to and high occupancy around those fixed points, which would correspond visually to long bars converging near those points. As one can see, the lengths and density of the bars are roughly uniform across both waking and sleep manifolds in a given session, Fig. 3d.

Separately but related, the change in angle is independent of the value of the angle itself, Fig. 3e (Fig. S8 for other animals). To obtain a quantifiable measure of occupancy along the ring and to gain statistical power from pooling across sessions and days from a single animal, we decode the angular states on the ring with a supervised decoder, and compute the density of decoded angles, Fig. 3f. The logarithm of the density of states along the ring - a direct estimate of the relative energy of these states - is flat on the scale of the variability across sessions. These results directly support property 4).

Finally, we examine the fluxes or net flows of states on versus off the manifold. The flux through a small region is the vector average over all the instantaneous trajectories that flow into and out of that region, Fig. 3g. For a continuous attractor not being driven by a directed input, we expect at most small net flows or fluxes along the manifold because of the uniform distribution of flows in all directions along the manifold (property 4), as well as because of the omnidirectional and unbiased nature of random kicks off the manifold; however, we expect large net fluxes for states that are not on the manifold because of biased flows of states returning to the manifold (property 3), precisely as seen in the data, where the fluxes are small on-manifold, but large for points off-manifold (Fig. 3g-i).

### Diffusive dynamics along manifold during REM

After obtaining a detailed *qualitative* picture of states and dynamics on and near the manifold, we combine theoretical predictions about dynamical trajectories on continuous attractor manifolds^40^ with our ability to perform time-resolved unsupervised decoding using SPUD (Fig. 4a-b) to gain a *quantitative* estimate of the nature and influence of noise on the circuit. Noise is an important consideration for integrator, memory, and representational circuits, because it determines the timescale and fidelity of information stored in the circuit.

**Figure 4:**
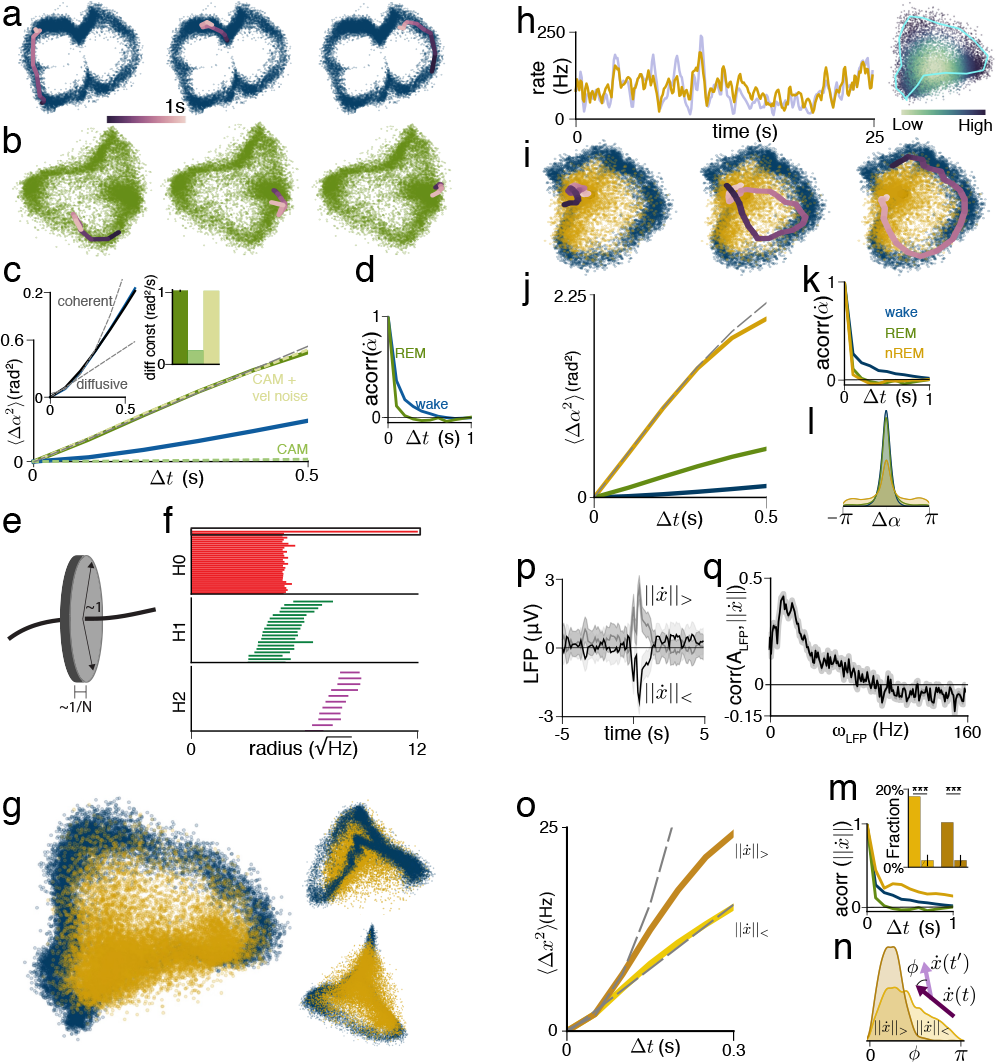
Diffusive dynamics during REM sleep, and higher-dimensional states and coherent dynamics during nREM sleep. (a) Manifold with sample waking trajectories at three different times. Each plot shows several consecutive points over a second and transitions between them. (b) As in (a) but for sample REM trajectories. (c) Plot of variance of REM angle update over time (solid green), with waking shown for comparison in blue. At short timescales, growth of variance is linear, with slope given by diffusion constant. Dashed green traces show continuous attractor model (CAM) with and without noisy velocity input. Left inset shows waking trace (blue: SPUD; black: measured head angle) on an expanded scale, highlighting supralinear increase at small times. Dashed lines in inset show pure diffusion and expected increase for velocity-driven dynamics. Bar plot shows diffusion constants for decoded angle (error bar is standard deviation from resampling with replacement), continuous attractor model without noisy velocity input, and continuous attractor model with noisy velocity input. (d) Autocorrelation of angular velocity. (e) Schematic showing that in high dimensions, independent noise is almost entirely directed off manifold and does not move the system much along the manifold. (f) Absence of persistent ring in Betti-1 barcode during nREM sleep (no long horizontal line; compare Figs. 2,3b). (g) Joint plot of nREM and waking manifolds using Isomap. nREM in mustard yellow; waking in dark blue (as before). Inset shows two alternative views. (h) Population firing rate during nREM decoded from distance to centroid (actual in light blue; decoded in yellow). Inset shows points colored by population firing rate. (i) nREM manifold with three sample trajectories. (j) Plot of variance of decoded nREM angle update over time (waking and REM shown for comparison). (k) Autocorrelation of angular velocity during nREM. (l) Distribution of changes in nREM angle over 500ms, with waking, REM angles in blue, green for comparison. (m) Autocorrelation of velocity on full manifold. Waking, REM traces from (k) shown for comparison. Inset shows fraction of time spent in consecutive low velocity and high velocity epochs (300ms duration each). Hatched bars show shuffled control (see SI S12). (n) Distribution of angles between successive high (dark) and low (light) velocities on full manifold (100ms separation). (o) Plot of squared change in position on full manifold for low and high velocity epochs. Quadratic and linear fits shown by dashed lines. (p) Average LFP trace conditioned on small (black) vs. large (grey) change in position, along with 95% confidence interval. (q) Correlation of total13change in position against LFP power in 1s bins, along with 95% confidence interval (see SI S12).

First, we validate that SPUD can sufficiently capture the fine-time-scale statistics of trajectories based on the available data by comparing time-lagged correlations of the unsupervised latent variable estimate during waking and against a “ground truth” of measured HD correlations, Fig.4c (blue and black traces in inset). Since HD updates during waking are correlated in time (Fig. 4d, blue trace and Supplementary Movie 3), the squared deflection in angle over time grows quadratically at short times (Fig. 4c, left inset).

During REM sleep, by contrast, angular updates are temporally uncorrelated but nevertheless local (SPUD result in Fig. 4d, green trace, and Supplementary Movie 4; the angle change histogram is small and unimodal), and the squared angular deflection grows linearly with time (Fig. 4c green curve). These two features — uncorrelated local updates together with a linear growth in squared deflection over time — are characteristic of an unbiased diffusive random walk, consistent with property 4) if the dynamics during sleep are noise-dominated and if the noise lacks temporally coherent structure^40^.

### Evidence of input aligned to manifold

To resolve the nature of the noise driving diffusivity during REM, we make, to our knowledge, the first quantitative comparison between empirically-observed diffusion in a neural circuit and theoretical predictions. The diffusion constant of REM dynamics in Fig. 4c is 1.1±0.04 *rad*^2^*/s* (0.52±0.03 and 1.3±0.06 for the other two animals; see Fig. S9). This diffusivity exceeds, by a factor of 20-50, the predicted value in a matched model (see^40^, SI S10 and Fig. S10), Fig. 4c, if the noise in the circuit is independent across neurons.

Independent per-neuron noise could arise from Poisson spike count variations within the circuit, or from a high-dimensional input that projects in a spatially uncorrelated way to the neurons. In either case, such high-dimensional noise is impotent in pushing the network state along the manifold, because noise of unit variance only has a projection of size 1*/N* along the manifold^4;40;41^, Fig. 4e. Increasing the amplitude of independent noise is not a solution: even 5x overdispersed noise does not account for the difference in diffusion constant, and independent noise of large magnitude destroys the low-dimensional states of the circuit (other factors that might potentially contribute to the gap between predicted versus observed diffusivity fall far short of accounting for the discrepancy, SI S10).

By contrast, a modest amount of noise in the form of fluctuations aligned to the nonlinear manifold, which is naturally interpreted as arriving in the velocity input to the circuit, has a much stronger effect^40;41^. Noise on the same order of magnitude as the velocity strengths required to produce HD-matched changes in represented angle in a model (standard deviation of 8.5 *rad/s*^1/2^, with temporal correlations of 20 ms or less; compare waking in Fig. S4), is sufficient to account for the measured diffusion, Fig. 4c. Noise aligned to the manifold additionally tends not to distort the activity states, and thus more purely moves the states along the manifold with at most small off-manifold effects, as apparently seen in the REM data. In sum, with high probability, REM diffusivity is driven by a low-dimensional noise injected into the circuit through an input aligned to the manifold, and thus likely arriving through the velocity pathway, with a magnitude consistent with the size of waking velocity inputs into the circuit (property 5).

These results demonstrate that even in high-level cognitive circuits for memory and integration, as previously established for low-level sensory circuits and sensorimotor pathways^42–45^, input noise or sensory precision rather than internal noise is the limiting factor for information fidelity.

### Inputs during nREM sleep disrupt ring manifold

Hippocampal circuits replay patterns of waking activity during nREM sleep^46–49^, and these events may play an important role in memory consolidation^50^. However, the HD circuit seems to lack replays or even coherent temporal dynamics^32;51^. On a more abstract level, nREM sleep is thought to disrupt the brain’s ability to maintain integrated representations^52^, but it is unclear what this means at a more granular level, for specific integrated representations like HD. A manifold-based approach reveals when previous population-level structure is modified and helps to understand the nature of the modification. In addition, if there is coherent dynamics in the restructured space, a manifold-based approach can help find it, even if the structure is not visible when states are projected into the old space.

The manifold in nREM does not preserve the ring structure of waking and REM states: it is higher-dimensional (persistent homology, contrast Fig. 4f with Fig. 2b, 3b; visualization, Fig. 4g and Supplementary Movie 5; also see correlation dimension estimates in Fig. S11), and only partially overlaps the waking/REM manifold (Fig. 4g).

The higher-dimensional nREM manifold encodes at least two latent variables. We decode an angle along the tangential (circular) dimension of the manifold using SPUD, and compare it with the outputs of two wake-trained supervised decoders that make different assumptions: a tuning-curve decoder and a population vector decoder (Fig. S12). The angles decoded by all three methods agree well (see Fig. S13 for another animal), except at low-activity states where the signal-to-noise ratio of the neural response is low.

A second latent variable obtained from the radial dimension of the manifold (based on distance to the manifold centroid) encodes the population firing rate, capturing slow, non-binary global rate fluctuations that characterize nREM sleep, Fig. 4h. Thus the nREM manifold is an amplitude-modulated version of the waking manifold^53;54^, forming a 2D conical surface (Fig. 4g and Fig. S11, where the cone is clearer for other animals). The circular boundary of the cone reaches toward the Wake ring, and the tip of the cone extends to the zero-activity state. These responses are well-modeled by the same circuit as for waking and REM dynamics - strong recurrent connectivity that supports the formation of an activity ring with local bump tuning of neurons - but modified so that the external input projecting globally to all neurons undergoes large amplitude fluctuations (Fig. S14). Unlike during REM, where external inputs permit the maintenance of the waking ring while driving states diffusively around it, during nREM the external inputs pull the states off the ring. Mechanistically, the difference is likely due to the loss, during nREM, of a discrete attractor dynamics that holds fixed the amplitude of background inputs across the large physiological shifts that occur between waking and REM.

The dynamics, like the states, are also higher-dimensional during nREM sleep, Fig. 4i. Examining the component of dynamics along only the angular dimension of the manifold, as would be done by a supervised decoder constructed from waking data, yields little temporal structure: nREM manifold trajectories projected onto the 1D waking ring rapidly decorrelate, Fig. 4j,k. The variance grows linearly with time, Fig. 4j, similar to the diffusive dynamics of angle during REM but with a much larger diffusion coefficient (~ 8 times REM), making it natural to interpret nREM dynamics as simply a faster version of the REM dynamics^32;55;56^ or otherwise unstructured dynamics^51^. However, fat tails in the histogram of state changes (Fig. 4l) suggest that state changes are not local and thus the dynamics are not actually diffusive.

Indeed, a different picture emerges for dynamics on the higher-dimensional manifold: there, we observe two distinct types of trajectories (partitioned by thresholds on the magnitude of displacement per unit time): periods of staying in a confined region of the nREM manifold and periods of large sweeps along large parts of it, Fig. 4i (and Supplementary Movie 6). The large sweeps are coherent: Large-displacement intervals occur successively over many intervals, or in long runs, compared to shuffled controls (Fig. 4m); motion along a direction tends to continue along that direction (Fig. 4n, dark histogram, showing the angle between successive displacement vectors over two 100ms intervals); and successive large displacement epochs show a quadratic growth (Fig. 4o, dark curve; 8x the speed of waking) in squared displacement over time, consistent with non-diffusive, directional motion. (In general, successive high-displacement epochs could simply look diffusive with a large diffusion coefficient; the quadratic growth is a clear symptom of coherent motion.)

The run lengths of successive small-displacement intervals are again overrepresented relative to shuffle controls (Fig. 4m), but unlike the sweeps, these small displacement epochs exhibit a linear growth in squared displacement over time (Fig. 4o, light curve), consistent with diffusive dynamics. The small-displacement intervals and sweeps during nREM each persist over longer timescales than are present in waking and REM dynamics; this persistence is responsible for the fat tail in the temporal autocorrelation dynamics of nREM states on the full manifold (compare 300 ms nREM correlations in Fig. 4m to 96 ms (waking, blue trace) and 38 ms (REM, green trace)).

Reproducing the dynamics of confined nREM trajectories in a circuit model (Fig. S14) requires temporally persistent low-activity states and slow amplitude modulation (0.1 Hz amplitude fluctuations thresholded to zero for about half the cycle, SI S13). However, sweeps cannot be explained simply by global fluctuations in population rate: most sweeps occur during high activity states, and the velocity during sweeps is not preferentially directed towards or away from the zero activity state (Fig. S15). The dynamics of sweeps require, in addition, temporally persistent or correlated velocity fluctuations (correlation time of 200 ms, the same as in the waking model and an order of magnitude slower than in the REM model).

To investigate the possible drivers of smooth sweeps, we correlate their occurance with the local field potential in ADn. The sweeps occur at times of transient increase in the local field potential amplitude, Fig. 4p, and specifically with transient upward fluctuations in the component of the LFP power around ~ 12 Hz, within the 7-15 Hz range for sleep spindles^57^), Fig. 4q. In turn, the occurrence of sleep spindles is correlated with the occurrence of sharp waves in the hippocampus^58^, raising the question of whether coherent sweeps in the HD circuit during sleep spindles have some relationship to replay and memory consolidation events occurring elsewhere in the brain^50^, including in the hippocampus^46–49^ and the neocortex^59;60^.

## Discussion

We have obtained a direct glimpse of the low-dimensional ring structure in the mammalian head-direction circuit, and by examining states across waking and sleep, have shown that the population dynamics are generated autonomously in the brain and are attractive. These direct observations, based on a manifold approach that reveals the full *N*-point correlations and dynamics of the circuit population response, complement and augment an elegant body of work which inferred low-dimensional structure from *pairwise correlations* in the vertebrate circuits for head direction^13;15;28–32^, oculomotor control^3;61^, prefrontal evidence accumulation^62^ and 2D spatial navigation^55;56;63^. Finally, the direct visualization of a clear one-dimensional ring in the activity states of the vertebrate HD circuit, where neurons may or may not be physically laid out in order of their activity profiles, provides a compelling parallel to the beautiful results on a topographically ordered ring recently discovered in the HD circuit of invertebrate nervous systems^18;21^.

We have sought to demonstrate that, as in the theoretical models of low-dimensional continuous attractor circuits, the natural way to understand neural circuits that represent low-dimensional variables is to examine the evolution of population states on a low-dimensional activity manifold. As we have shown, this approach allows for the observation of attractive dynamics, which would have been difficult to demonstrate otherwise, and for the quantification of dynamics on the manifold. It also allows for the discovery of coherent dynamics when the states are altered, as we saw in nREM sleep. Finally, it allows for comparison with theoretical models, whose key predictions are at the level of structured population dynamics.

The unsupervised extraction of the encoded variable and neural tuning curves from manifold characterization provided an estimate that outperformed measured head angle in accounting for neural spikes. The manifold approach to latent variable discovery and decoding is useful whenever the encoded variable or some subset of the encoded variables is unknown: this is often the case in cognitive systems, but can also be true for sensory and motor systems when they are modulated by top-down and other inputs; and it is usually the case during sleep^50^. It will also be useful in examining how structured states and dynamics emerge in neural circuits during development^64^, plasticity, or learning.

In the case of HD cells, individual neural tuning curves are sparse, local functions on the manifold. However, SPUD and related manifold discovery methods^33;34;65;66^ (D. Tank, personal communication; Y. Ziv, personal communication) can be used for unsupervised decoding and tuning curve discovery when tuning curves are highly non-sparse and non-local. Manifold methods can also be fruitfully applied to topologically non-trivial manifolds of higher dimension^b^, such as a toroidal structure produced by simulated grid cells (SI S3; ~35 cells are sufficient to visualize 2-dimensional toroidal structure if the tuning curves are not too narrow, Fig. S3).

In sum, manifold-level analyses can enable fully unsupervised discovery and decoding of brain states and dynamics, and the quantification of collective dynamics on and off the manifold can give insight into circuit mechanism. We believe that manifold learning and related techniques^33;34;67;68^ will be essential for extracting information from large datasets, representing the future of neural decoding.

## Methods Summary

Information on the data set and preprocessing are in Supplementary Information S1. Information on the methods we use to extract and parametrize low-dimensional structure are in Supplementary Information S2 and S4. Details of the waking decoding are in S5, for REM decoding in S7-9, and nREM decoding in S11, 12. The model construction is described in S6, S10 and S13. Data have been previously reported in^32^ and are available on CRCNS: http://crcns.org/data-sets/thalamus/th-1. Code is available on request.

## Footnotes

a The manifold is sufficiently convoluted that it does not occupy a very low-dimensional linear subspace, thus linear embedding methods are of limited use and can create artefactual self-intersections in the manifold projection. Nonlinear embedding methods ^37–39^ are less prone to these problems.

b The number of cells required to characterize a manifold of dimension *Dm* grows with *Dm* in the exponent; this scaling is less a property of a specific method than of the intrinsic complexity of characterizing higher-dimensional structures, commonly called the curse of dimensionality.

## References

[1] Amari, S.-I. Dynamics of pattern formation in lateral-inhibition type neural fields. Biol. Cybern. 27, 77–87 (1977).

[2] Hopfield, J. J. Neural networks and physical systems with emergent collective computational abilities. Proc. Natl. Acad. Sci. U. S. A. 79, 2554–8 (1982).

[3] Seung, H. S. How the brain keeps the eyes still. Proc. Natl. Acad. Sci. U. S. A. 93, 13339–13344 (1996).

[4] Zhang, K. Representation of spatial orientation by the intrinsic dynamics of the head-direction cell ensemble: a theory. J. Neurosci. 15, 2112–2126 (1996).

[5] Seung, H. S., Lee, D. D., Reis, B. Y. & Tank, D. W. Stability of the memory of eye position in a recurrent network of conductance-based model neurons. Neuron 26, 259–271 (2000).

[6] Deneve, S., Latham, P. E. & Pouget, A. Efficient computation and cue integration with noisy population codes. Nat. Neurosci. 4, 826 (2001).

[7] Burak, Y. & Fiete, I. R. Accurate path integration in continuous attractor network models of grid cells. PLoS Comput. Biol. 5, e1000291 (2009).

[8] Shenoy, K. V., Sahani, M. & Churchland, M. M. Cortical control of arm movements: a dynamical systems perspective. Annu. Rev. Neurosci. 36, 337–359 (2013).

[9] Mazor, O. & Laurent, G. Transient dynamics versus fixed points in odor representations by locust antennal lobe projection neurons. Neuron 48, 661–673 (2005).

[10] Mante, V., Sussillo, D., Shenoy, K. V. & Newsome, W. T. Context-dependent computation by recurrent dynamics in prefrontal cortex. Nature 503, 78–84 (2013).

[11] Saha, D. et al. A spatiotemporal coding mechanism for background-invariant odor recognition. Nat. Neurosci. 16, 1830–1839 (2013).

[12] Gallego, J. A., Perich, M. G., Miller, L. E. & Solla, S. A. Neural manifolds for the control of movement. Neuron 94, 978–984 (2017).

[13] Ranck, J. B. Head direction cells in the deep cell layer of dorsolateral pre-subiculum in freely moving rats. In Buzsaki, G. & Vanderwolf, C. (eds.) Electrical Activity of Archicortex, 217–220 (Akademiai Kiado, 1985).

[14] Taube, J. S., Muller, R. U. & Ranck, J. B. Head-direction cells recorded from the post-subiculum in freely moving rats. I. Description and quantitative analysis. J. Neurosci. 10, 420–435 (1990).

[15] Taube, J. S., Muller, R. U. & Ranck, J. B. Head-direction cells recorded from the postsubiculum in freely moving rats. II. Effects of environmental manipulations. J. Neurosci. 10, 436–447 (1990).

[16] Sharp, P. E., Blair, H. T. & Cho, J. The anatomical and computational basis of the rat head-direction cell signal. Trends Neurosci. 24, 289–294 (2001).

[17] Finkelstein, A. et al. Three-dimensional head-direction coding in the bat brain. Nature 517, 159–164 (2015).

[18] Seelig, J. D. & Jayaraman, V. Neural dynamics for landmark orientation and angular path integration. Nature 521, 186–191 (2015).

[19] Turner-Evans, D. et al. Angular velocity integration in a fly heading circuit. eLife 6, e23496 (2017).

[20] Green, J. et al. A neural circuit architecture for angular integration in Drosophila. Nature 546, 101–106 (2017).

[21] Kim, S. S., Rouault, H., Druckmann, S. & Jayaraman, V. Ring attractor dynamics in the Drosophila central brain. Science 356, 849–853 (2017).

[22] Skaggs, W. E., Knierim, J. J., Kudrimoti, H. S. & McNaughton, B. L. A model of the neural basis of the rat’s sense of direction. In Advances in Neural Information Processing Systems 7 (NIPS), 173–180 (1995).

[23] Blair, H. T. Simulation of a thalamocortical circuit for computing directional heading in the rat. In Advances in Neural Information Processing Systems 8 (NIPS), 152–158 (1996).

[24] Redish, A. D., Elga, A. N. & Touretzky, D. S. A coupled attractor model of the rodent head direction system. Netw. Comput. Neural Syst. 7, 671–685 (1996).

[25] Sharp, P. E., Blair, H. T. & Brown, M. Neural network modeling of the hippocampal formation spatial signals and their possible role in navigation: a modular approach. Hippocampus 6, 720–734 (1996).

[26] Aronov, D., Nevers, R. & Tank, D. W. Mapping of a non-spatial dimension by the hippocampal–entorhinal circuit. Nature 543, 719–722 (2017).

[27] Killian, N. J., Jutras, M. J. & Buffalo, E. A. A map of visual space in the primate entorhinal cortex. Nature 491, 761–764 (2012).

[28] Mizumori, S. & Williams, J. Directionally selective mnemonic properties of neurons in the lateral dorsal nucleus of the thalamus of rats. J. Neurosci. 13, 4015–4028 (1993).

[29] Taube, J. S. & Burton, H. L. Head direction cell activity monitored in a novel environment and during a cue conflict situation. J. Neurophysiology 74, 1953–1971 (1995).

[30] Knierim, J. J., Kudrimoti, H. S. & McNaughton, B. L. Interactions between idiothetic cues and external landmarks in the control of place cells and head direction cells. J. Neurophysiol. 80, 425–446 (1998).

[31] Yoganarasimha, D., Yu, X. & Knierim, J. J. Head direction cell representations maintain internal coherence during conflicting proximal and distal cue rotations: comparison with hippocampal place cells. J. Neurosci. 26, 622–631 (2006).

[32] Peyrache, A., Lacroix, M. M., Petersen, P. C. & Buzsáki, G. Internally organized mechanisms of the head direction sense. Nat. Neurosci. 18, 569–575 (2015).

[33] Chaudhuri, R., Gercek, B., Pandey, B. & Fiete, I. Unsupervised latent variable extraction from neural data to characterize processing across states. In Computational and Systems Neuroscience (CoSyNe), I–56 (Salt Lake City, UT, 2017).

[34] Chaudhuri, R., Gercek, B., Pandey, B. & Fiete, I. Unsupervised latent variable extraction from neural data to characterize processing across states. In Annual Meeting of Society for Neuroscience (SfN), 346.13 (Washington, DC, 2017).

[35] Ghrist, R. Barcodes: The persistent topology of data. Bull. Amer. Math. Soc. 45, 61–75 (2008).

[36] Grassberger, P. & Procaccia, I. Measuring the strangeness of strange attractors. Physica D 9, 189–208 (1983).

[37] Tenenbaum, J. B., De Silva, V. & Langford, J. C. A global geometric framework for nonlinear dimensionality reduction. Science 290, 2319–2323 (2000).

[38] Roweis, S. T. & Saul, L. K. Nonlinear dimensionality reduction by locally linear embedding. Science 290, 2323–2326 (2000).

[39] Kingma, D. P. & Welling, M. Auto-encoding variational bayes. Proceedings of the International Conference on Learning Representations (ICLR) (2014).

[40] Burak, Y. & Fiete, I. R. Fundamental limits on persistent activity in networks of noisy neurons. Proc. Natl. Acad. Sci. U.S.A. 109, 17645–50 (2012).

[41] Moreno-Bote, R. et al. Information-limiting correlations. Nat. Neurosci. 17, 1410–1417 (2014).

[42] Bialek, W. Physical limits to sensation and perception. Annu. Rev. Biophys. Biophys. Chem. 16, 455–478 (1987).

[43] Bialek, W. & Owen, W. G. Temporal filtering in retinal bipolar cells. elements of an optimal computation? Biophys. J. 58, 1227–1233 (1990).

[44] Osborne, L. C., Lisberger, S. G. & Bialek, W. A sensory source for motor variation. Nature 437, 412–416 (2005).

[45] Pitkow, X. & Meister, M. Decorrelation and efficient coding by retinal ganglion cells. Nat. Neurosci. 15, 628–635 (2012).

[46] Pavlides, C. & Winson, J. Influences of hippocampal place cell firing in the awake state on the activity of these cells during subsequent sleep episodes. J. Neurosci. 9, 2907–2918 (1989).

[47] Wilson, M. A. & McNaughton, B. L. Reactivation of hippocampal ensemble memories during sleep. Science 265, 676–679 (1994).

[48] Skaggs, W. E. & McNaughton, B. L. Replay of neuronal firing sequences in rat hippocampus during sleep following spatial experience. Science 271, 1870–1873 (1996).

[49] Lee, A. K. & Wilson, M. A. Memory of sequential experience in the hippocampus during slow wave sleep. Neuron 36, 1183–1194 (2002).

[50] Diekelmann, S. & Born, J. The memory function of sleep. Nat. Rev. Neurosci. 11, 114–126 (2010).

[51] Brandon, M. P., Bogaard, A. R., Andrews, C. M. & Hasselmo, M. E. Head direction cells in the postsubiculum do not show replay of prior waking sequences during sleep. Hippocampus 22, 604–618 (2012).

[52] Massimini, M. et al. Breakdown of cortical effective connectivity during sleep. Science 309, 2228–2232 (2005).

[53] Goris, R. L., Movshon, J. A. & Simoncelli, E. P. Partitioning neuronal variability. Nat. Neurosci. 17, 858–865 (2014).

[54] McCormick, D. A., McGinley, M. J. & Salkoff, D. B. Brain state dependent activity in the cortex and thalamus. Curr. Opin. Neurobiol. 31, 133–140 (2015).

[55] Gardner, R. J., Lu, L., Wernle, T., Moser, M.-B. & Moser, E. I. Correlation structure of grid cells is preserved during sleep. bioRxiv 198499 (2017).

[56] Trettel, S. G., Trimper, J. B., Hwaun, E., Fiete, I. R. & Colgin, L. L. Grid cell co-activity patterns during sleep reflect spatial overlap of grid fields during active behaviors. bioRxiv 198671 (2017).

[57] Lüthi, A. Sleep spindles: where they come from, what they do. Neuroscientist 20, 243–256 (2014).

[58] Siapas, A. G. & Wilson, M. A. Coordinated interactions between hippocampal ripples and cortical spindles during slow-wave sleep. Neuron 21, 1123–1128 (1998).

[59] Ji, D. & Wilson, M. A. Coordinated memory replay in the visual cortex and hippocampus during sleep. Nat. Neurosci. 10, 100–107 (2007).

[60] Rothschild, G., Eban, E. & Frank, L. M. A cortical-hippocampal-cortical loop of information processing during memory consolidation. Nat. Neurosci. 20, 251259 (2017).

[61] Aksay, E., Gamkrelidze, G., Seung, H., Baker, R. & Tank, D. In vivo intracellular recording and perturbation of persistent activity in a neural integrator. Nat. Neurosci. 4, 184–193 (2001).

[62] Wimmer, K., Nykamp, D. Q., Constantinidis, C. & Compte, A. Bump attractor dynamics in prefrontal cortex explains behavioral precision in spatial working memory. Nat. Neurosci. 17, 431–439 (2014).

[63] Yoon, K. et al. Specific evidence of low-dimensional continuous attractor dynamics in grid cells. Nat. Neurosci. 16, 1077–84 (2013).

[64] Bassett, J. P., Wills, T. J. & Cacucci, F. Self-organised attractor dynamics in the developing head direction circuit. Curr. Biol. 28, 609–615 (2018).

[65] Park, M. et al. Bayesian Manifold Learning: the Locally Linear Latent Variable Model. In Advances in Neural Information Processing Systems 28 (NIPS), 154–162 (2015).

[66] Rybakken, E., Baas, N. & Dunn, B. Decoding of neural data using cohomological learning. bioRxiv 222331 (2018).

[67] Linderman, S. W., Johnson, M. J., Wilson, M. A. & Chen, Z. A Bayesian nonparametric approach for uncovering rat hippocampal population codes during spatial navigation. J. Neurosci. Methods 263, 36–47 (2016).

[68] Wu, A., Roy, N. G., Keeley, S. & Pillow, J. W. Gaussian process based nonlinear latent structure discovery in multivariate spike train data. In Advances in Neural Information Processing Systems 30 (NIPS), 3496–3505 (2017).

