## Supplementary Information for "The population dynamics of a canonical cognitive circuit"

#### S1 Data and preprocessing

Data have been previously reported in[1] and are available on CRCNS: http://crcns.org/data-sets/thalamus/th-1. The data set contains recordings from the anterodorsal thalamic nucleus (ADn) of 7 mice. We show manifolds from all 7 mice and decoded angles from the 3 mice with root-mean-square decoding error of $< 0.5$ rad (see decoding section for details).

Sessions that allow good decoding contain 9-50 neurons, and the number of neurons that show good head direction tuning ranges from 8-30. We do not preselect neurons in any way. Our decoding quality improves rapidly with the number of neurons, suggesting that SPUD is likely to perform well on larger datasets with low-dimensional structure.

For some of the mice, the data also contains recordings from the postsubiculum. Including data from the postsubiculum allows for better decoding of waking head direction. However the manifold from the postsubiculum encodes other behavioral variables and is strongly modulated by arousal state (not shown), and so we do not include these data.

In the main text we show Mouse 28, session 140313 for Figs. 2 and 4, and Mouse 25, session 140130 for Fig. 3.

##### Data binning and kernel smoothing

We first convert the series of spike times for the set of $N$ simultaneously recorded neurons into time-varying counts or rates. To do this, we do one of the following: 1) Construct non-overlapping bins of width $\Delta t$ across the recording time $[0, T]$, then per neuron and per bin, replace the spikes within each bin with the spike count. The result is an $N \times (T/\Delta t)$ matrix $\mathbf{C}$ of spike counts $C_{it}$ for neuron $i$ and time-step $t$. 2) Convolve the spike train in time with a Gaussian kernel (of standard deviation $\sigma$ and unit area) to yield a matrix of smoothly varying spike rate estimates across neurons and time.

For most of our analyses (except those specifically mentioned in the following sentences), we use the spike train convolved with a Gaussian kernel of standard deviation $\sigma = 100$ms to estimate the time-varying rates, and construct point clouds from these rates sampled at 100ms intervals. For the persistent homology measures we use non-overlapping bins of width 1s, and for the diffusion plot in Figure 4o we use $\sigma = 50$ ms to ensure that we do not miss fine timescale structure (results are very similar with $\sigma = 100$ms). Results are similar across choices of bin sizes/kernel widths, though getting good estimates of fine timescale dynamics requires bins/kernels with widths on the order of 100ms, and persistent homology performs best with bins of 1s.

##### Supervised tuning curve extraction and decoding

Here we describe the conventional, supervised method used to estimate neural tuning curves and perform decoding. We also refer to this method as the tuning curve (TC) decoder. To extract the supervised tuning curve of a cell, we use head angle data (measured directly from

LEDs placed on the animal's head[1]) and calculate the mean response of the cell for each angular bin. The tuning curve of the $i$-th cell, $f_i(\theta)$, is:

$$f_i(\theta) = \frac{\text{Number of spikes fired by cell } i \text{ around angle } \theta}{\text{Time spent by animal around angle } \theta}.$$

We perform supervised decoding of head angle by maximum likelihood estimation from the neural data and the tuning curves, under the model that at angle $\theta$, neuron $i$ responds with a number $C_i$ of Poisson distributed spikes with rate given by its tuning curve $f_i(\theta)$, and that the responses of different neurons are independent, conditioned on their individual tuning curves. Thus,

$$\hat{\theta}_t = \arg\max_{\theta} P(\{C_{it}\}_{i=1,\cdots,N}|\theta) = \arg\max_{\theta} \prod_{i=1}^{N} Poiss(C_{it}; f_i(\theta)\Delta t)$$

For constructing tuning curves to the SPUD decoded angle, we use the same procedure, except replacing the measured angle $\theta$ with the SPUD angle $\alpha$.

In both cases we use 30 bins between 0 and $2\pi$ to construct the tuning curves.

##### Variance stabilization/Gaussianization

After binning but before applying our unsupervised decoding methods, we replace spike counts with their square roots. Under the assumption that spike counts are generated from a Poisson process (with mean given by the instantaneous firing rate), this transformation stabilizes the variance, ensuring that variance does not depend on the mean[2]. We find this transformation dramatically improves the quality of our visualization and decoding.

### S2 Low-dimensional structure in the head-direction system

#### S2.1 Visualization and noise-reduction via dimensionality reduction

Before fitting the manifold or applying topological methods, we first reduce the large ($N$-dimensional) ambient dimension by re-embedding the data into a smaller, but still relatively high-dimensional *embedding space* of dimension $D_e$ ($D_m \ll D_e \ll N$, where as before $D_m$ is the intrinsic manifold dimension; note that according to results from Dimension Theory[3], the embedding dimension must satisfy $D_e \geq 2D_m + 1$ to avoid constructing artefactual self-intersections of the manifold – e.g. to avoid mapping a convoluted ring manifold into a figure-eight).

Nonlinear dimensionality reduction or embedding into a sufficiently high-dimensional embedding space ($D_e \geq 2D_m + 1$) irons out some convolutions in the manifold while preserving its topology. The reduced dimension speeds up spline fitting (a practical concern), and ironing out the convolutions appears to yield modest improvements in SPUD decoding

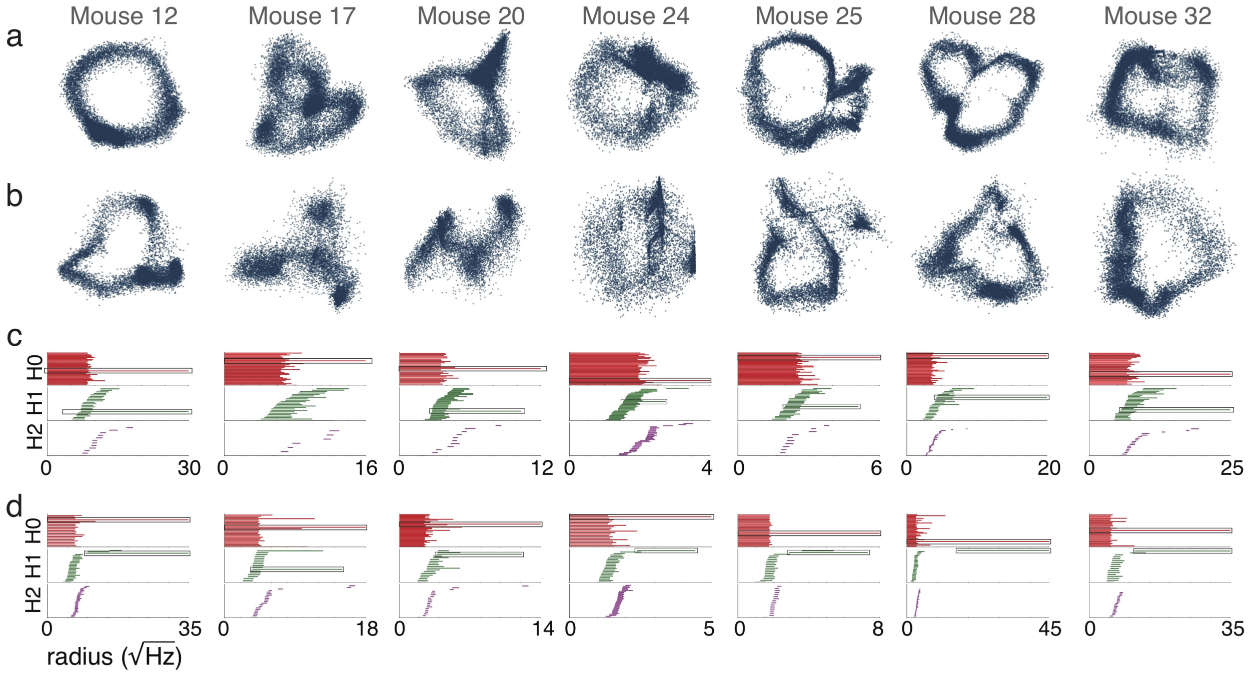


Figure S1: **Low-dimensional structure in the head direction circuit.** Each column shows results from a different animal in the study of[1]. (a) Waking manifold visualized using Isomap[4]. (b) Waking manifold visualized using a Variational AutoEncoder[5;6]. (c) Betti barcodes on data without outlier removal. Top row shows Betti-0 (number of connected components); second row shows Betti-1 (number of rings); third row shows Betti-2 (number of three-dimensional holes or voids). (d) As in (c) but for data with outliers removed (see S2.2 on nt-TDA).

performance (Fig. S4). As noted in the main text, the results for decoding are not sensitive to the choice of embedding dimension (also see section on wake decoding).

In what follows we use the nonlinear dimensionality reduction method Isomap[4] as a preprocessing step throughout. We set the number of neighbors to be 5 (higher values work well too), and embed into 3-20 dimensions (3 for visualization and before decoding; 10 before applying the topological methods below; a range between 3-20 to characterize decoding for the plot in Fig. S4a). For the joint visualizations of data across states (Fig. 3c, 4g) we concatenate equal amounts of data from the two states (determined by the state of shorter duration) and run Isomap on this combined data. Note that Isomap is a nonlinear method, and thus distances in the embedding space are not a simple global rescaling of distances in the full firing rate space.

In Fig. S1a we show the underlying ring manifold for all 7 animals in the data of[1] using Isomap.

We also compared Isomap to other dimensionality reduction methods: a Variational Autoencoder (VAE)[5;6], shown in Fig. S1b; Locally Linear Embedding (LLE)[7]; and Principal Components Analysis (PCA).

We found that Isomap typically performed better than the VAE, which was unable to extract clear rings from several of the sessions, possibly reflecting its sensitivity to the limited-data regime. One potential advantage of the VAE method over the other nonlinear dimensionality reduction methods is that it allows an estimate of manifold dimension based on the number of active units in the bottleneck layer (the smallest hidden layer between the encoder and decoder that the VAE uses to reconstruct the data). However, we find that the VAE uses two active units, thus estimating the manifold dimension as 2. This reflects the minimum number of dimensions required to embed a ring manifold in Euclidean space, rather than the intrinsic manifold dimension of 1.

Isomap also performed better than Locally Linear Embedding (LLE)[7]. LLE was often able to reveal a ring (and when it did it could produce rings with less scatter than the other methods). However, LLE often failed for small binwidths ($\sim 100$ ms) and for these reasons we do not show results from LLE.

Finally, linear dimensionality reduction methods like PCA are sometimes able to find a ring in the ADn dataset for waking states, though they perform worse than nonlinear dimensionality reduction. PCA can distort the manifold so that certain angles are over- or under-represented, and can force intersections in the low-dimensional projection when none are present in the high-dimensional space (see Fig. S2a for examples). These problems can be understood from the sparse structure of the individual cell responses, as is the case when the data are generated from narrow tuning curves to a low-dimensional variable: As shown in Fig. S2b, if there are $N$ cells with narrow tuning curves and relatively small overlaps, the 1D coding manifold narrowly hugs every axis in the ambient space, despite its intrinsic low-dimensionality. In such cases, the manifold does not lie on any low-dimensional linear subspace of the ambient space. For this reason, linear subspace methods like PCA cannot preserve the manifold structure in a reduced-dimensional projection.

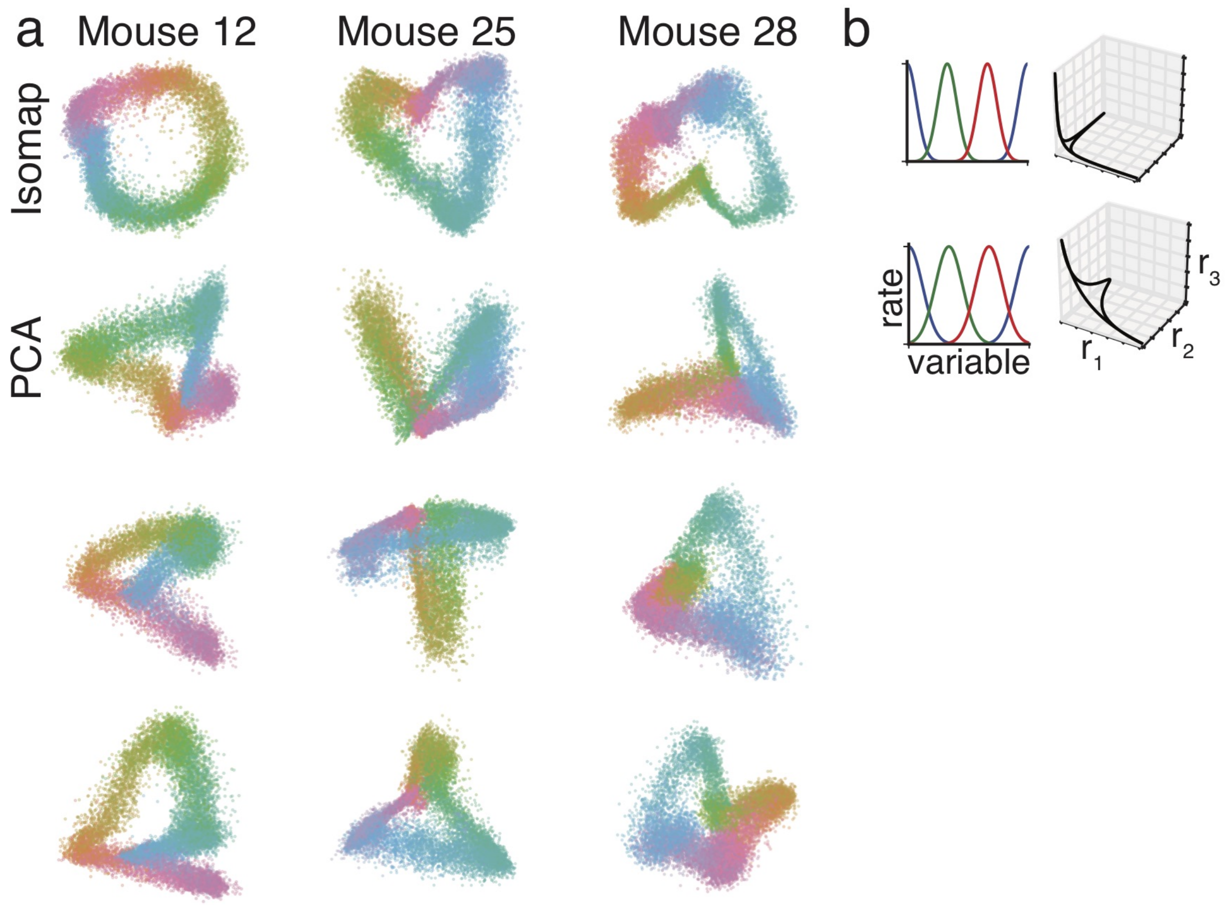


Figure S2: **Distortion of manifold by linear methods.** (a) Each column shows results from a different animal, highlighting a manifold that is distorted by PCA. Top row shows Isomap projection, while next 3 rows show PCA projection (three different views). Manifolds are colored by measured head angle. (b) Left column: tuning curves for three model neurons tuned to a one-dimensional variable (synthetic data). Right column: corresponding manifold in state space. Top shows narrow tuning curves; bottom shows wide tuning curves. Manifold is one-dimensional but increasingly nonlinear for narrower tuning curves.

#### S2.2 Persistent homology

We use and assess methods that characterize, through the measure of *persistent homology*, the topology of the manifold assumed to underlie the data[8;9].

Given a point cloud of data in a metric space, and some spatial scale (distance) $d$, these methods first construct a simplicial complex by connecting all points within a distance $d$, Fig. 1b. A 0-simplex is a point, a 1-simplex is a connected pair of points (i.e., an edge), a 2-simplex is a triangle, a 3-simplex is a tetrahedron, and so on. If the point cloud is drawn from an underlying continuous space, then the topological properties of the simplicial complex will reflect the topological properties of the underlying space, such as the number of distinct pieces, closed 1-dimensional loops, 2-dimensional voids enclosed by the space, etc.

For a given value of $d$, the corresponding simplicial complex reflects the properties of the underlying space at that particular spatial scale. The next step is to consider a range of values of $d$ and to examine how features vary with $d$. For example, Fig. 1b shows two 1-dimensional loops that are each visible for a particular range of values of $d$. Persistent homology looks for topological features of a data set that persist over a range of spatial scales, guided by the intuition that such persistent features are likely to reflect important aspects of the data rather than noise.

The set of simplicial complexes at each thresholded level are assigned *Betti numbers* that characterize their topological characteristics (for instance, a simplicial complex with a ring-like topology has a non-zero Betti-1 number). The existence of different non-zero Betti numbers is traced across different threshold levels, to generate the so-called *Betti bar code* (Fig. 1b, far right). Non-zero Betti numbers that exist across many levels are called persistent; a persistent number signifies that the corresponding topological feature (a ring in the case of a persistent Betti-1 number) is a robust and important property of the data.

Methods based on persistent homology are appealing because they allow data to be simultaneously examined at multiple length scales, and are finding increasing use in neuroscience[10;11]. Nevertheless, these methods do not typically contain a noise model or measures of statistical significance and thus can be hard to interpret on very noisy data. Relatedly, they are highly sensitive to outliers: a noisy point in the data can easily split one persistent ring into two. Thus more data is not necessarily better when applying these methods. Such a splitting into multiple rings can be seen in some of the REM sessions (Fig. S7), though the lack of a significant Betti-2 suggests that these multiple rings do not reflect the presence of a torus in the data. Extending persistent homology methods to a more probabilistic setting is an active area of research[12;13], especially deserving of study in neural systems, where responses are highly variable[14;15], but at the same time, there are sometimes reasonable statistical models for the generation of such variability[16;17]. As described below, we use a simple thresholding method to exclude outliers. This greatly improves the extraction of features during REM sleep. During waking the features are already very robust, and are slightly improved by the thresholding method.

To compute Betti barcodes for the data we first apply Isomap to the binned spike counts (bin size of 1s) to reduce it to 10 dimensions (or use all neurons if the number of neurons is $\leq 10$). We then use the package Ripser[18] to generate the Betti-0, -1 and -2 barcodes, reflecting the Betti numbers at different spatial scales, and plot the most persistent features in the figures (i.e., we do not plot bars below some minimum length, chosen independently for each figure). We also repeat this analysis after excluding outliers by (a) first considering a neighborhood around each point with radius defined by the 1st percentile of the pairwise distance distribution, and (b) then removing all points whose number of neighbors lie in the bottom 20th percentile of the distribution of number of neighbors across points (referred to as nt-TDA in text and figure captions). We show both sets of results in all figures that follow.

As seen in Fig. S1c,d the Betti-1 barcodes for the data show a persistent ring structure during waking.

#### S2.3 Correlation dimension to compare manifolds across states

The *correlation dimension*[19] quantifies the intrinsic dimension of a manifold locally, by counting how the number of manifold points ($M_r$) contained in a state-space ball centered at some point on the manifold expands as the ball expands. The number will grow as $r^{D_m}$ for a manifold of intrinsic dimension $D_m$ if $r$ is the ball radius (Fig. 1c). The slope of $\log(M_r(r))$ provides an estimate of dimension. In general, this estimate is an upper-bound on dimensionality: If the sampled neural states are noisy, with some independent noise per neuron, the manifold will be "fluffy" at the scale of the noise standard deviation $\sigma_M$, and for small ball radii ($r < \sigma_M$) the estimated manifold dimension will be high. If the manifold is curved, folding upon itself so that distant regions of the manifold come within a distance $R$ in the state space, large balls (with $r > R$) will include more than one part of the manifold and the dimension estimate will again exceed the true dimension. Thus, if $\sigma_M \sim R$, the estimate all values of $r$ will be a systematic overestimate of the manifold dimension. However, if $\sigma_M \ll R$, the function $M_r(r)$ will exhibit a pure power-law region at $r$ intermediate between $\sigma_M$ and $R$, with the power providing an accurate estimate of the local intrinsic dimension.

To examine whether correlation dimension can estimate manifold dimension accurately, we generate synthetic data from a population of 20 neurons responding to a $k$-dimensional stimulus ($k$ from 1 to 5). Stimulus values are drawn from $[0, 1]^k$. Each neuron in our sample has a tuning curve with center $\mu_i$ (distributed in $[0, 1]^k$), width $\sigma_i$ (uniform between 0.2 and 0.4) and peak firing rate $R_i = 20$ Hz. Given a stimulus value $x$, neuron $i$ generates spike counts $C_i \sim Poiss\left(R_i e^{-(x-\mu_i)^2/2\sigma_i^2}\right)$. Thus, the population responses form a $k$-dimensional manifold, with position on the manifold corresponding to the stimulus $x$. We then embed the data into a 10-dimensional space using Isomap (identical to preprocessing for persistent homology), compute the number of points within a distance of $r$, as $r$ is varied, and fit a straight line to this curve in log-log space.

Our analyses of this synthetic data show that in the presence of neurally-plausible levels of noise (i.e., Poisson-like variability), correlation dimension is a poor estimator of absolute manifold dimension, but the estimated correlated dimension reflects the relative ordering of actual dimension: if correlation dimension of network A is higher than that of network B, then so is the dimension of the variable that network A encodes (data not shown). Thus we use correlation dimension to compare the dimension of the representation within an area across states (Fig. S11d). As with synthetic data, we first embed the data into a 10-dimensional space using Isomap, or use all neurons if the number of cells is $\leq 10$. We compute the number of points within a distance of $r$, as $r$ is varied, and fit a straight line to this curve in log-log space using the birth and death of the persistent homology ring to choose the fit range.

Correlation dimension does not contain a noise model. Indeed the separation between signal and noise is not well-defined without further information, because noise is indistinguishable from a high-dimensional manifold. As we discussed for persistent homology methods, responses in neural systems are highly variable[14;15], but there are often reasonable statistical models for this variability[16;17], and thus it will be interesting to extend correlation dimension measures to neural data by explicitly attempting to model this variability.

### S3 Manifold structure in simulated grid cell data

To illustrate how higher-dimensional manifold structure might be inferred from data and assess the number of cells that might be required for a topologically non-trivial 2D manifold, we generate synthetic data from model grid cells[20], whose states are periodic in two dimensions. Grid cells with a common period and orientation (thus from a single module) are expected to exhibit a low-dimensional state-space manifold[21], specifically the states should lie on a 2D torus.

The synthetic grid cell tuning curves are generated from a bivariate von Mises distribution as

$$f_i(\phi, \psi) = e^{(5\cos(\phi-\mu)+5\cos(\psi-\nu)-\cos(\phi-\mu-\psi+\nu))}$$

(For simplicity, we have generated square lattices, but the results will be essentially the same for a triangular lattice.) Here $f_i(\phi, \psi)$ is the firing rate of the $i$th cell at location $(\phi,\ \psi)$. The cell has receptive field center $(\mu_i,\ \nu_i)$ and the peak firing rate is normalized to 50 Hz. Spikes are generated from these tuning curves as a Poisson process.

A torus has one connected component (Betti-0 of 1), two rings or one-dimensional holes (Betti-1 of 2) and one void or two-dimensional hole (Betti-2 of 1). We find that $\sim 35$ cells, if the preferred phases are sufficiently well spread-out, are sufficient to visualize the toroidal manifold, Fig. S3a. Similarly, if there are more than $\sim 35$ cells, persistent homology can recover these Betti numbers with good fidelity, Fig. S3b.

### S4 Spline fit and decoding

##### Spline fit to manifold

We fit the manifolds by piecewise linear curves. A curve is specified by a set of $K$ knots, with locations $x_i$. The knots are ordered, and the $i$-th segment of a curve is a straight line between the $i$th and $i+1$th knot. To fit these curves given a number of knots $K$, we first use k-means to identify $K$ clusters in the data and set the centers of these clusters to be the initial knot locations. We then iteratively update these knot locations to minimize the distance of the data points from the piecewise linear curve formed by these knots. Note that this method could be extended to add smoothness constraints at the knots.

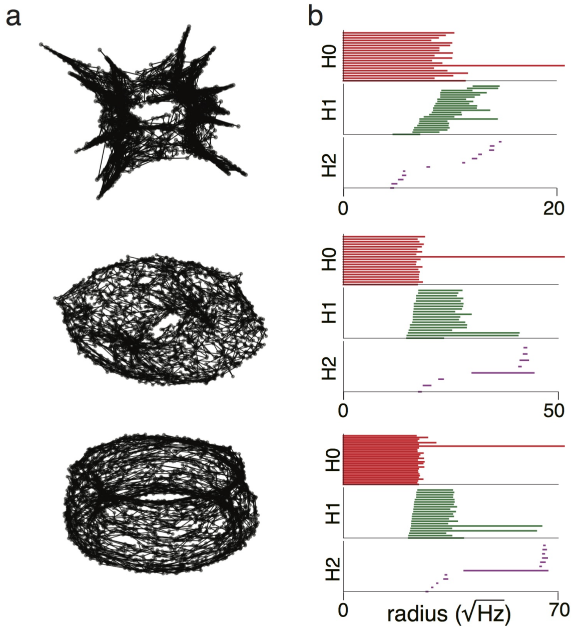


Figure S3: **Toroidal manifold in simulated grid cell activity** (a) Manifold extracted from synthetic grid cell spiking data (IID Poisson spikes based on grid-like activation of multiple neurons from the same simulated module – in other words, the simulated cells share a common spatial tuning period and orientation, and with a uniform spread of 2D spatial phase). Top to bottom shows $N = 16, 36, 64$ (b) Corresponding Betti barcodes for data shown in (a): there are 1-2 significant non-zero Betti-1 numbers throughout (1 in the top plot as expected for a ring because of insufficient data; 2 in the latter two, as expected for a torus), and a significant non-zero Betti-2 number (in the bottom two, as expected for a torus).

##### Parameterization of spline and decoding

We parameterize points on the manifold by distance along the curve from some arbitrary origin, with distances rescaled to lie between 0 and $2\pi$ for comparison to actual head angle. The length of each line segment is taken to be proportional to its distance in the embedding space (we also tried assigning each line segment an equal length so that if there are $K$ such line segments, then each has length $2\pi/K$ and results were similar though slightly worse).

In the waking state, where there is a ground truth measured head angle, we shift the global origin and choose the orientation around the curve to match observed head angle, but made no other modifications (e.g., we did not rescale the coordinate differently in different parts of the ring). During sleep, when comparing to a tuning curve decoder we perform a similar shift and choice of orientation dictated by the tuning curve decoded angle.

Points are decoded by mapping them to the nearest point on the manifold, based on Euclidean norm in the embedding space, and reading off the parameter value there.

#### Model parameters

The only parameters in the model—all of which are fit to the data—are the coordinates of the $K$ anchor points used in the spline and the estimated manifold dimension $D_m$ (a total of $KD_e + 1$ numbers, independent of the number of neurons $N$ and the duration of the time-series; increasing the data volume through recording duration or neuron numbers only increases the signal-to-noise ratio of this fixed number of parameters, thus the performance of the method will improve). There are four scalar hyperparameters: $dt$, the temporal bin size to convert spike times into time-varying counts; $K$, the number of anchors to use in the spline; and $D_e$ and $M$, the embedding dimension and number of neighbors respectively for the noise-reducing embedding step (the parameter $M$ is specific to the nonlinear dimensionality reduction method we used; some other methods do not require it). The most sensitive parameter is $D_m$; the method is relatively insensitive to the detailed values of the hyperparameters.

For the decoding shown in the main text we used data smoothed with a Gaussian kernel of $dt = 100$ ms standard deviation, and set $K = 12$. We primarily fit in $D_e = 3$ dimensions (after doing Isomap with $M = 5$ neighbors, described in previous section), but show higher dimensional fits in Fig. S4. We choose the change in coordinates along a line segment of the spline to be proportional to its distance in the embedding space.

### S5 Decoding during waking

#### Dependence on embedding dimension and on quality of ring

For the results shown in the main text, we performed the spline fit and decoding after first embedding the data in 3 dimensions. In Fig. S4a, we show that we can fit and parameterize the manifold in a much higher-dimensional embedding space, at the cost of a small growth in squared error with embedding dimension, consistent with a random noise-based decline in the signal-to-noise ratio for less dimensionally-reduced embeddings.

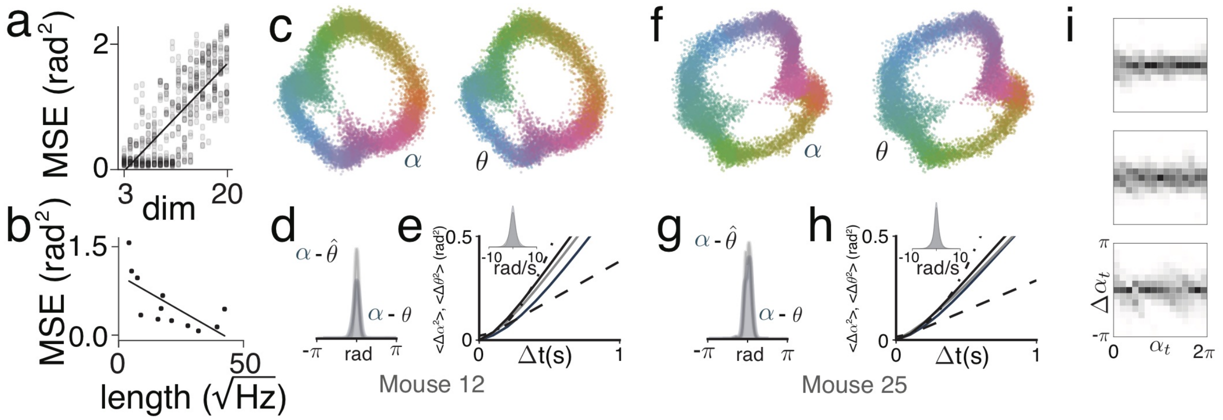


Figure S4: **Wake decoding across animals** (a) Linear increase in squared "error" (defined as difference between unsupervised latent variable estimate and measured HD) as a function of embedding dimension $D_e$ ($r = 0.78$). (b) Decrease in decoding error for more persistent ring feature ($r = -0.69$). Persistence measured as the length of the longest Betti-1 feature (shown in Fig. S1). (c) Waking manifold for Mouse 12 colored by SPUD (left) and actual (right) head angle. (d) Difference of SPUD angle from a tuning curve decoded angle (gray) and actual head angle (blue). (e) Plot of the average squared change in angle for different time separations, highlighting supralinear increase at small times. Dashed lines show pure diffusion and expected increase for velocity-driven dynamics. Blue, gray, black show SPUD, TC and actual angle respectively. Inset shows distribution of measured (black) and SPUD (blue) velocities. (f-h) Same as (a-c) for another animal, Mouse 25. (i) Conditional distribution of changes in angle (over 500ms) given starting angle, shown for 3 animals. Top to bottom: Mouse 28 (other statistics shown in main text), Mouse 12 and Mouse 25).

We also find that the decoding performance is strongly correlated with the length of the persistent feature extracted by persistent homology (i.e., the length of the longest Betti-1 bar shown in Fig. S1b): a more persistent feature yields better decoding, Fig. S4b.

#### Decoding performance across animals

In the main text we show decoding for Mouse 28. In Fig. S4c-h we show that similar results hold for the other two animals. SPUD is able to extract head angle and fine timescale dynamics with accuracy comparable to a supervised decoder, and the dynamics are compatible with a velocity-driven representation.

#### Homogeneous dynamics

The dynamics on the manifold during waking are homogeneous, meaning that the distribution of changes to the latent variable does not depend on the value of the latent variable (Fig. S4i), as would be expected for a representation that continually integrates an angular velocity input.

#### Variance explained

To compute the variance explained (as in Fig. 2i), we considered the spike counts extracted in 100 ms bins. If the spike counts of the $i$th neuron are $C_i$, then the variance explained by the variable X is:

$$\mathrm{Var}_{\mathrm{exp,X}} = \mathrm{Var}\left[\mathbb{E}(C_i|X)\right] + \mathbb{E}\left[\phi \mathbb{E}(C_i|X)\right]. \tag{1}$$

Here $X$ can be the measured head angle or one of several decoded head angles, including the supervised tuning curve estimate and various unsupervised latent variable estimates. We bin the variable X to lie in one of 30 bins between 0 and $2\pi$. For the Poisson model, $\phi = 1$. For the overdispersed model, we estimate $\phi$ as $\min_X \mathrm{Var}(C_i|X)/\mathbb{E}(C_i|X)$ where, as before, $X$ is the variable we condition upon. Note that for a true overdispersed process taking the minimum is likely to underestimate the overdispersion but we choose this in order to be conservative.

In all cases (except for measured head angle), we divided the data into a training set (80% of data) and a test set (remaining 20%). We used the training set to fit the manifold or construct the tuning curves, and evaluated the explained variance on the test set. The explained variances plotted are the averages across all cells.

In Fig. 2i, we evaluated significance by computing the number of cells which were better explained by the unsupervised latent variable estimate than the measured angle, and comparing this to a null model where both measured and estimated angles explained the data equally well (i.e., binomial distribution with $p = 0.5$; two-sided test).

In Fig. S5a we compare the variance explained by $\theta$ and $\alpha$ to the variance explained by the supervised tuning curve decoder (gray) and to two other unsupervised estimates.

For the SPUD-tuning-curve estimate (Fig. S5a, brown bars), we construct tuning curves of each neuron to the unsupervised latent variable estimate, and then use these tuning curves to do decoding. This additional step allows some averaging away of noise and thus allows for slightly better decoding.

For the leave-one-out estimate (Fig. S5a, red bars), we estimate the latent variable without using the activity of one neuron, and then examine how much of that neuron's variance is explained by the latent variable. Define the set of firing rate vectors $r_{\setminus i}(t)$ to be the activity of all neurons excluding neuron $i$. Then, to generate the unsupervised latent variable estimate to use for neuron $i$ at time $t$, we take the vector $r_{\setminus i}(t)$, find its 5 nearest neighbors (in Euclidean distance) in the training set, and estimate the latent variable as the average of the unsupervised latent variable estimate for these neighbors. We then compute the variance explained for neuron $i$, as described above. We repeat this analysis for each neuron and average the resulting variance explained. This decreased the variance explained slightly for Mouse 12 and Mouse 28 ($2-3\%$) and had a larger effect on Mouse 25. Mouse 25 has 10 recorded neurons while the other two have $> 20$ cells. Given this small number of neurons, removing a single neuron degrades the unsupervised latent variable estimate, causing performance to drop.

In Fig. 2k we plotted the covariance between the spike counts along with the average covariance after conditioning on either measured or SPUD angle.

For Fig. 2j, we generated synthetic data using the tuning curves to the unsupervised latent variable estimate. Given a decoded latent variable $\alpha$, we generate a spike count for neuron $i$ from a normal distribution with mean $\mathbb{E}(C_i|\alpha)$ and variance $\mathrm{Var}(C_i|\alpha)$. Generating

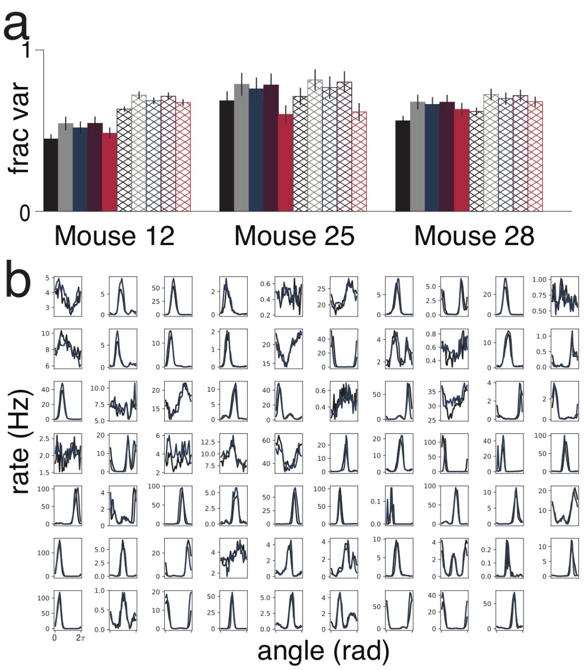


Figure S5: **Spiking variance and tuning curves explained by decoding** (a) Average fraction of variance explained by (left to right): measured angle, supervised estimate from tuning curve decoder (gray), unsupervised latent variable estimate (blue), SPUD-tuning-curve-estimate (brown), leave-one-out estimate (red). Solid bars show Poisson model and hatched bars show overdispersed model. (b) Plot of tuning curves to measured angle ($\theta$, black) and unsupervised latent variable estimate ($\alpha$, blue). Variance of tuning curves explained by $\alpha$ is $71\% \pm 2.8\%$.

counts this way assumes that neural firing is overdispersed but that neurons are independent given $\alpha$, thus explicitly removing any additional structure in the population.

### S6 Continuous attractor model reproduces waking dynamics

We perform similar analyses on a continuous attractor model of the head direction system as we do on the data, and show that the model reproduces the low-dimensional structure and dynamics empirically observed. We use a modified version of the model from [22]. Each neuron in the model is tuned to a preferred head angle and receives inhibitory input from a ring of surrounding neurons that are tuned to other head angles (Fig. S6a). This inhibition is small for neurons with similar tuning and increases as the difference in tuning increases. The network as a whole receives nonspecific feedforward excitation. With appropriate parameters, the population response is a localized bump of activity whose location on the ring corresponds to the represented head angle.

To couple the model to velocity input, we consider two rings of the form described above (Fig. S6b). Connectivity within one ring is shifted a few degrees in the clockwise direction and connectivity in the other ring is shifted in the counterclockwise direction. As a consequence of this asymmetric connectivity, neurons in one ring drive the population activity clockwise along the ring while neurons in the other ring drive the population activity counterclockwise. The two rings are coupled together with symmetric non-shifted inputs. If the activity of the neurons in the two rings is balanced, then the population response will remain in place on the ring. We model velocity input as a change in gain to neural activity in these two rings: thus clockwise head movement increases the activity of neurons in the clockwise ring and suppresses the activity of neurons in the other ring; consequently, the population response is driven along the rings in the appropriate direction.

Each neuron is described by the equation

$$\tau \frac{ds_i}{dt} = -s_i + \sum_{\alpha} \delta(t - t_i^{\alpha}), \tag{2}$$

where $s_i$ is the synaptic activation of the $i$th neuron, $\tau$ is the synaptic time-constant, and $t_i^{\alpha}$ are the spike times. The $i$th neuron fires spikes with rate given by

$$r_i = \phi(g_i) = \phi\left(\sum_j W_{ij} s_j + b_i\right). \tag{3}$$

Here $\phi$ is the f-I curve, $g_i$ is the synaptic input, $W_{ij}$ is the connection strength from neuron $j$ to neuron $i$, and $b_i$ is a non-specific background input, here taken to be the same across all neurons. Spikes are generated according to a Poisson distribution with this rate.

Each neuron within a ring has a preferred angle, given by $\theta_i = 2\pi i/N$, where $N$ is the total number of neurons in each ring and $i$ is the index of the neuron. The connectivity between two neurons with preferred angles $\theta_i$ and $\theta_j$ is $W_{ij} = w(\theta_i - \theta_j + \Delta)$ if neuron $j$ is in the clockwise ring and is $W_{ij} = w(\theta_i - \theta_j - \Delta)$ if neuron $j$ is in the counterclockwise ring. The kernel $w$ is defined by,

$$w(\theta) = \exp\left[k_1(\cos(\theta) - 1)\right] - \exp\left[k_2(\cos(\theta) - 1)\right]. \tag{4}$$

A non-zero velocity signal, $v$, rescales the gain of input to each ring. Thus $g_i = (1 \pm \alpha v_i)\left(\sum_j W_{ij} s_j + b_i\right)$, where the sign is positive for the counterclockwise ring and negative for the clockwise ring.

Parameters are: Synaptic time-constant $\tau = 10$ ms; f-I curve $\phi(x) = 10e^x/\tau$; background input $b_i = -2.85$; connectivity shift $\Delta = 2$ rad; connectivity kernel parameters $k_1 = 1$, $k_2 = 0.3$; gain of velocity input $\alpha = 0.1$; number of neurons $N = 1000$ for simulations of waking/nREM and 4000 for REM (number is unimportant except when characterizing the intrinsic diffusion of the activity bump/angle representation, which depends inversely on number).

In Fig. S6c, we show sample trajectories when this model is driven by a velocity input that is correlated over time, as when the animal is awake and exploring the environment. To generate these visualizations we subsample 25 neurons from the network, bin the spikes

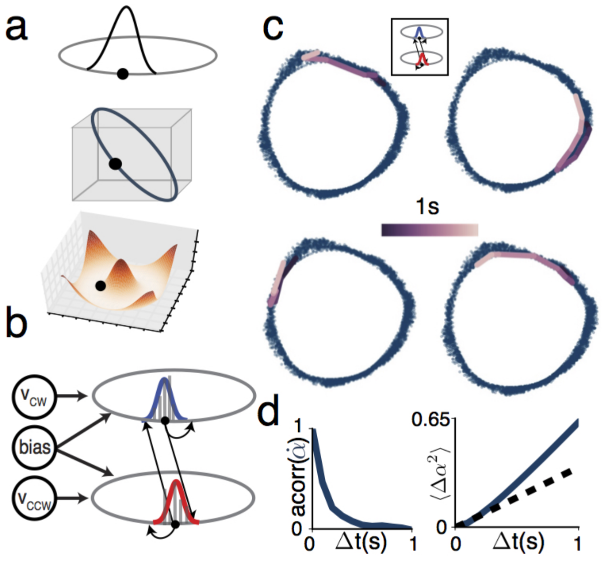


Figure S6: **Wake model** (a) Schematic of a continuous ring attractor model. The model consists of a ring of neurons, which maintain a localized bump of activity whose position corresponds to the encoded variable (top). The population activity forms a one-dimensional ring in neural state space (middle). The dynamics can be understood as movement downhill on an energy landscape (bottom). (b) Two connected ring models, one driving the attractor bump clockwise (top) and the other driving the bump counterclockwise (bottom). (c) Same analysis as in Fig. 4a applied to model simulations, showing four sample trajectories. (d) Left: Autocorrelation of successive change in model angle. Right: Diffusion plot of change in model angle.

in 100 ms bins, and then apply Isomap to the square root of these counts (as for the real data). As in the trajectories of Fig. 4a, typical trajectories in the model consist of smooth, relatively slow sweeps on the manifold. Changes in angle are autocorrelated on a timescale of hundreds of milliseconds (Fig. S6d, left panel). Moreover, a diffusion plot showing the variance of the distribution of changes in elapsed angle over time reveals quadratic behavior at short timescales (Fig. S6d, right panel), as expected for a state driven by a slowly-varying velocity. Thus, the dynamics we see in the data when the animal is awake are compatible with a continuous attractor that integrates a slowly-varying angular velocity input to update head angle.

The velocity input was generated from a zero-mean Ornstein-Uhlenbeck process, with standard deviation 1 $rad/s^{1/2}$, and correlation time 200ms.

### S7 One-dimensional ring manifold during REM

In Fig. S7a we show that the REM manifold is identical to the waking manifolds across all animals. Moreover, persistent homology finds persistent ring structure, Fig. S7b,c though the ring sometimes fragments into multiple rings due to outliers (note that the absence of

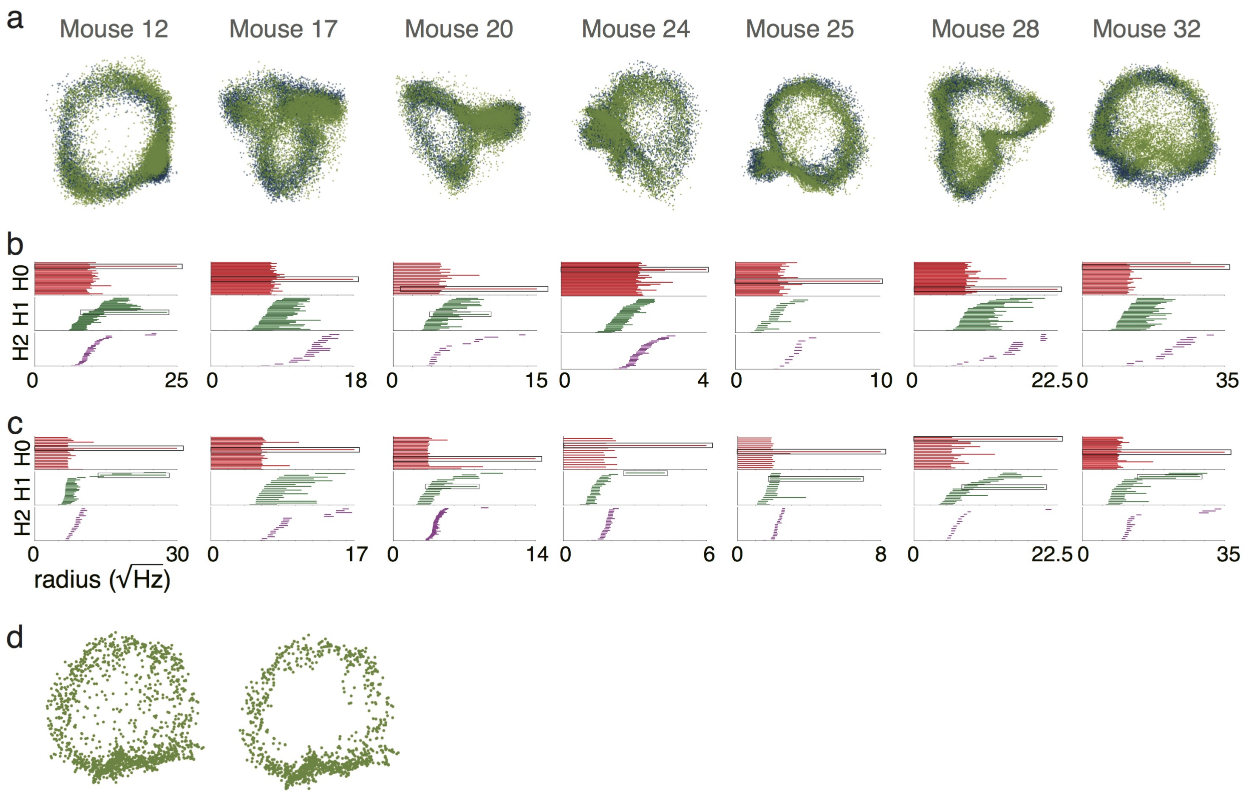


Figure S7: **REM manifold across animals** (a) Joint visualization of waking (blue) and REM (green) manifolds across all animals. (b) Betti-0, -1 and -2 barcodes for REM manifold using data without outlier removal. (c) As in (b) but for data with outliers removed (see S2.2 on nt-TDA). (d) Manifold from Mouse 25 before (left) and after (right) removal of outliers for nt-TDA. Note that TDA uses 1s bins so there are fewer points than in panel (a).

Betti-2 means that the manifold is not a torus).

### S8 Flows and manifold occupancy during REM sleep

In Fig. 3e, we plot the mean change in the extracted latent variable on a single session as a function of its value, to show that mean changes are small compared to the standard deviation and thus that dynamics are homogeneous and diffusive. In Fig. S8a, d we show that the same is true for Mouse 12 and Mouse 28.

For the occupancy plot in Fig. 3f, we use a tuning curve decoder to combine sessions from Mouse 25 to get a distribution of occupancies across all sessions, including those sessions on which there were not enough neurons to extract and parameterize a ring manifold (this is the only place where we rely on supervised decoding), and plot the mean and standard deviation of occupancy. Note that decoded occupancy could vary over recording sessions for a variety of reasons, including the particular sample of neurons that were recorded. In Fig.

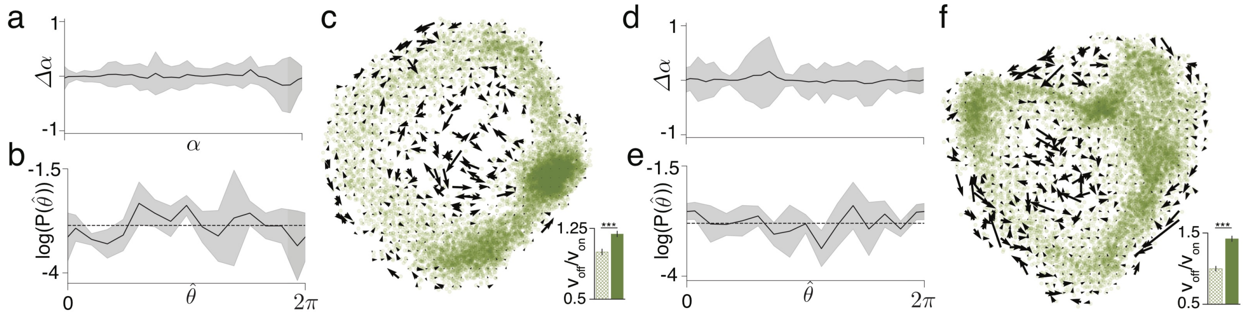

Figure S8: **REM occupancy and flows** (a) Single session mean and standard deviation of change in decoded angle as a function of angle for Mouse 12. (b) Angle occupancy mean and standard deviation (from tuning curve decoder) across sessions. (c) Flux on and off manifold. Flux is larger off manifold (Inset shows shuffled control (hatched bar); $p < 10^{-6}$). (d-f) As in (a-c) but for Mouse 28. $p < 10^{-6}$.

S8b, e we show the occupancy for the other two mice we consider.

For the flow fields in Fig. 3g, we considered a two-dimensional embedding of the REM manifold, grouped points into 2025 (i.e, 45 x 45) bins and averaged together the velocity vectors of all points in a bin to reveal that the average velocity vector is larger off manifold than on. In Fig. S8c, f we show these average velocity vectors for the other two mice we consider.

To quantify the difference between on and off manifold points, we divide the bins into on and off-manifold bins based on the 50th percentile of the distance to the fitted spline (i.e., $<$ 50th percentile is on manifold), and compare the average norm of the velocity vector on and off manifold (bar plot in Fig. 3i and Fig. S8c, f; error bars show standard deviation of ratios resampled with replacement). There are fewer points off manifold than on and thus these results might have resulted from averaging together a smaller number of random vectors off manifold. To control for this, we shuffle the assignment of velocity vectors to points and recompute the ratio of on to off manifold velocity vectors to generate a null distribution (shown as control in bar plots of Fig. 3i and Fig. S8c, f; error bars show standard deviation across permuted samples). For all 3 animals, the average velocity vector is significantly larger off manifold than on ($p < 10^{-6}$), with significance computed by comparing the observed ratio to the distribution of shuffled ratios.

### S9 Dynamics during REM sleep

As during waking, during REM sleep changes to the latent variable do not depend on the value of the latent variable (Fig. S9a), reflecting homogeneous diffusion along the manifold. To compute the diffusion constants we fit a straight line to the first 200 ms of the squared change in decoded angle against time.

Applying SPUD to data from ADn for the other two animals yields similar results as for Mouse28, Fig. S9b-g: the ring manifolds are present, though noisier, and the dynamics

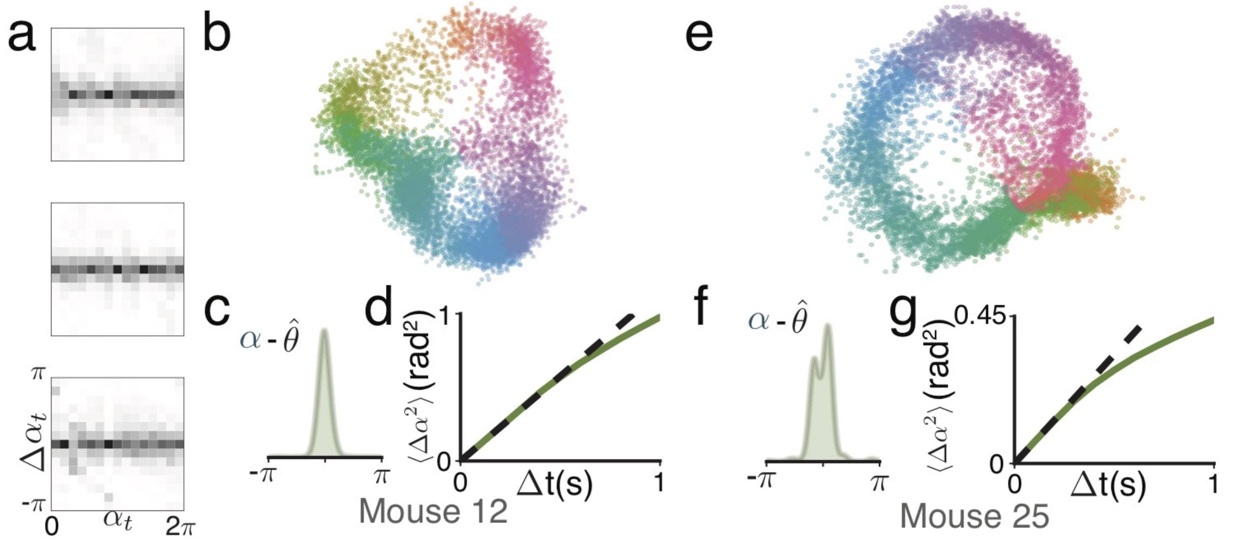


Figure S9: **REM decoding across animals** (a) Conditional distribution of changes in angle (over 500ms) given starting angle, shown for 3 animals. Top to bottom: Mouse 28 (other statistics shown in main text), Mouse 12 and Mouse 25. (b) REM manifold colored by SPUD angle for Mouse 12. (c) Difference of SPUD angle from a tuning curve decoded angle. (d) Plot of the average squared change in angle for different time separations, along with linear (diffusive) fit. (e-g) Same as (b-d) for Mouse 25.

are diffusive, as predicted by a continuous attractor model. As for Mouse 28, the diffusion constants are too large to be explained by Poisson-like fluctuations in neural spiking, and instead suggest noisy input through the velocity pathway (see discussion in main text).

### S10 REM dynamics replicated in a continuous attractor model

It is possible to model the dynamics and dynamical statistics of REM states using the same continuous-attractor ring model in Fig. S6, with the modification that there is either no velocity input, or that the input from the velocity channel is random and temporally uncorrelated. Both these models agree qualitatively with the data, but only one of them is capable of providing a quantitative match.

In the model without velocity input, independent Poisson spiking noise across neurons within the network pushes the population states around the 1D ring of stable states. The model exhibits low-dimensional trajectories similar to those seen during REM sleep (Fig. S10a), and successive angle updates are uncorrelated, except at very short timescales (Fig. S10b, left panel). As predicted by theoretical analyses, the represented angle performs a noise-driven random walk (resulting from integrating internal noise fluctuations instead of a coherent velocity input), and the squared deviation in angle over a small time interval $\Delta t$ grows linearly with $\Delta t$ (Fig. 4c and Fig. S10b, right panel), just as observed during REM sleep.

However, the rate of diffusion in the data is much larger than in the model for parameters roughly matched to quantities in ADn ($N = 4000$ neurons in model, according to an estimate of the number of head direction cells in mouse ADn; peak neural firing rates of 20 Hz; independent, identically distributed Poisson fluctuations in neural spike counts), Fig. 4c. The discrepancy is 40-fold. A contributing factor to the discrepancy may be an over-optimistic theoretical estimate of the average peak firing rates (by interpreting recorded cells as "typical" when they are an experimentally selected, highly modulated subset), but closing the gap with this factor would require that the typical peak firing rate across all neurons is 40 times smaller than estimated; further, at this low rate, the model would not exhibit ring attractor states. Similarly, increasing the magnitude of independent noise in the HD circuit's neural activities cannot account for the discrepancy: even if spikes were overdispersed with a variance equal to five times the mean firing rate (modeled by replacing the Poisson spike generation by a normal distribution with variance five times the mean), which greatly exceeds the modest overdispersion of 2.4 we determine from the ADn data (analysis not shown). Further increasing the overdispersion in the model leads to death of the activity bumps. As explained in the main text, neural spiking noise is high-dimensional and produces only a tiny projection along the angular coding dimension of the network, in inverse proportion to network size. Thus, even large amplitudes of independent or high-dimensional noise are not able to produce much change in the represented angle (this noise tolerance is the main hypothesized reason for why the brain constructs low-dimensional attractors), and are more likely to disrupt the existence of attractor states than to move them along the manifold.

Imperfections in the recurrent weights underlying bump states would produce a deviation from perfect continuous attractor dynamics and result in discrete stable states, to which the internal states are attracted. Such drifts of state could be fast and thus consistent with a larger squared deviation in represented angle over time than possible with unbiased noise; however, these drifts would be unidirectional (correlated) in time, and would thus produce a quadratic diffusivity curve, rather than the curves observed in the REM data (Fig. 4c). In sum, diffusive drift in the HD system during REM sleep is unlikely to be explained by sampling biases, independent neural fluctuations or weight imperfections within the HD circuit.

If noise were projected along a particular direction in the state space, specifically along the 1D angle-representing ring, then modest amounts of this low-dimensional noise would be able to cause large changes in bump position. For temporally uncorrelated and unbiased noise of this type, the changes in bump position in the network would additionally be diffusive. In an integrator circuit, the only low-dimensional projection along the nonlinear coding manifold is the velocity input. We add input fluctuations along the velocity pathway to the ring model (filtered random Gaussian inputs, with short autocorrelation time of 20 ms and standard deviation fit to match the observed diffusion constant). For the traces shown in Fig. 4c and S10b, the the standard deviation providing a good fit to the observed diffusivity of REM dynamics is $8.5 rad/s^{1/2}$. This fluctuation amplitude is the same order of magnitude as the amplitude of velocity inputs required to produce the measured waking displacements in the model.

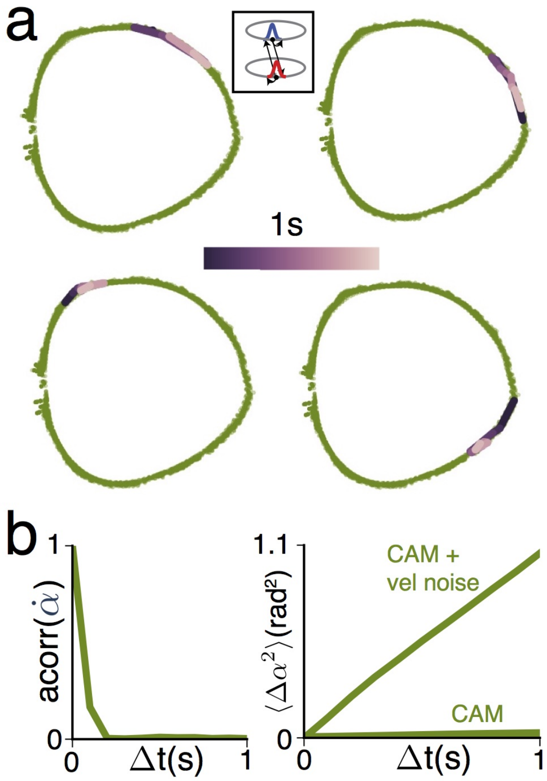


Figure S10: **REM states and dynamics replicated in a continuous attractor model** (a) Sample trajectories from model, visualized as in Fig. 4a. (b) Left: Autocorrelation of angle updates in model. Right: Diffusion plot for model simulated with and without velocity input noise (same as model traces from Fig. 4c).

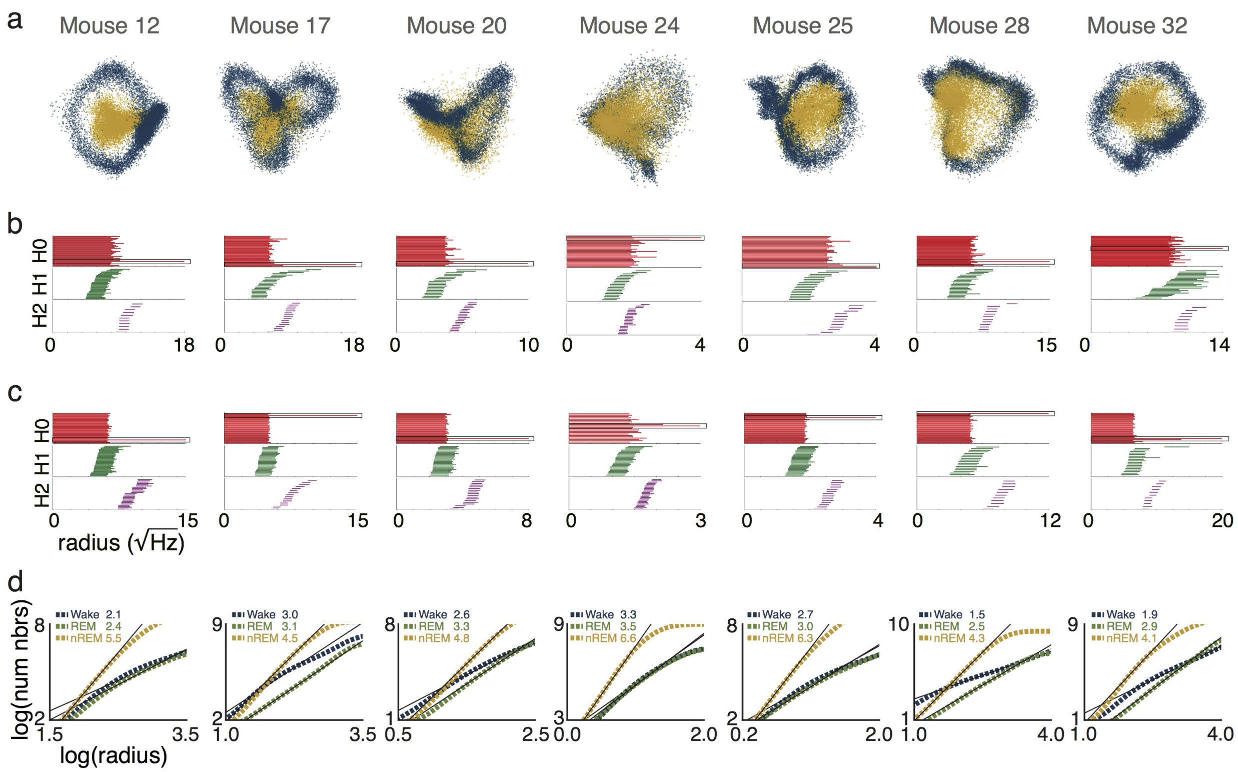


Figure S11: **nREM manifold across animals** (a) Joint visualization of waking (blue) and nREM (mustard yellow) manifolds across all animals. (b) Betti-0, -1 and -2 barcodes for nREM manifold with TDA (no outlier removal as in nt-TDA). (c) As in (b) but for data with outliers removed (see S2.2 on nt-TDA) (d) Scaling of number of neighbors against radius (the slope of the central portion of these curves is the local correlation dimension of the manifold) across states. The nREM manifolds are consistently higher-dimensional than waking/REM manifolds, as shown by the steepness of the nREM curves and the greater slope of fitted lines for nREM.

### S11 Loss of ring manifold during nREM

As with Mouse 28 in the main text, across all animals the nREM manifold is not a ring and is higher-dimensional, as can be seen by direct visualization, Fig. S11a, the absence of persistent rings (no significant feature in the Betti-1 barcode), Fig. S11b,c, and the higher estimated correlation dimension, Fig. S11d (see S2.3 for description of method).

The degree of modulation in ADn during nREM varies by animal: for example, the nREM manifold of Mouse 25 significantly overlaps the waking manifold, while the nREM and wake manifolds are separable in Mouse 12 . However, our qualitative conclusion holds across animals: during nREM, population states are pulled off of the waking manifold and towards the zero activity state.

### S12 Decoding during nREM sleep

In the main text, we argue that ADn population activity during nREM sleep still reflects an angular variable despite distortions in the manifold induced by global fluctuations. Of course simply projecting the population representation to a ring manifold will always result in an angular variable. We thus support our conclusion in two independent ways, Fig. S12.

First, we decoded nREM states from ADn using a tuning curve decoder (derived from waking data), binned the decoded angle, and showed the average population activity for each decoded angle bin, Fig. S12a. The panels in Fig. S12a show the typical pattern of nREM population activity that corresponds to each decoded angle (mustard curves). Note that the population activity takes the form of a population-level activity bump that moves systematically across angle bins. Moreover, the population average response curves are very similar in shape across states (results similar for distributions of activities rather than means; not shown), with nREM activity lower in amplitude than the other two states. The largest discrepancies are in low activity bins (Bins 5 and 12 for this animal), reflecting the poor decoding of low activity states during nREM. Low-activity states could result either from an angle that is poorly represented in our sample of neurons or from global activity fluctuations at other angles. The tuning curve decoder will map all low activity states to angles with poor coverage in our sample, and do so with high posterior probabilitiy (because a low-activity state resulting from a global amplitude fluctation at a different angle looks very similar to the normal state for a specific angle with poor neural coverage in our sample). Consequently, these angles appear to have high occupancy during nREM sleep.

Second, we compare SPUD to two wake-trained decoders that make different assumptions: a tuning-curve decoder (described previously) and a population vector (PV) decoder that attempts to correct for global fluctuations. To construct this decoder we consider the set of vectors from the origin to each point on the waking/REM manifold and, given a new population vector (i.e., during nREM sleep), we assigned it the manifold coordinate corresponding to the waking vector that subtends the smallest angle with it. Thus, the PV decoder uses only the direction of the population firing rate vector and not its magnitude.

SPUD, the TC decoder and the PV decoder all agree very well despite making quite different assumptions, Fig. S12b, except at very low activity states where, as previously argued, the decoding problem is ill-posed.

For SPUD-based firing rate decoding in Fig. 4h, we compute the distance of points on the nREM manifold to the manifold centroid, and plot the best linear predictor of population firing rate as a function of this distance.

To classify trajectories on the full manifold as sweeps, we set a threshold of the 60th percentile of the speed and look for 300 ms epochs (here 6 50ms time bins; results are similar for 3 100ms time bins) when the speed is above this threshold. For confined trajectories we do the same but extract trajectories that remain below the 40th percentile of speed. For the controls, we shuffle the velocities over time and apply the same analysis. Error bars for the controls show standard deviation across samples, and we calculate significance using the shuffled distribution.

For the analyses involving the local field potential (LFP), we estimate the LFP at a shank as the median of the LFP recorded on each channel, and then average these estimates across shanks (results are similar across shanks). For Fig. 4p, we consider changes in

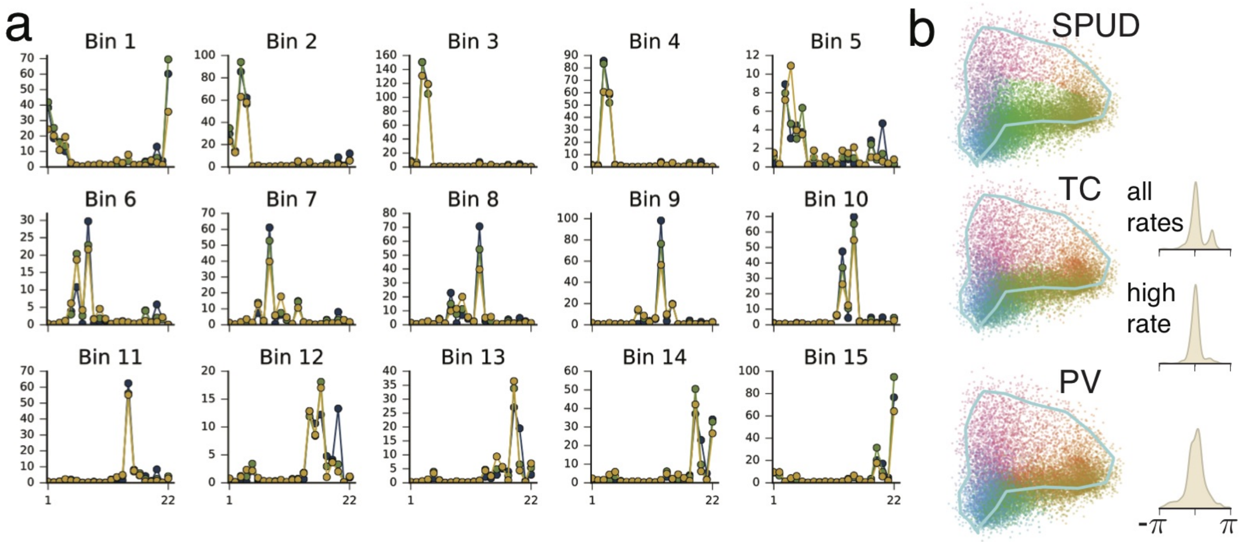


Figure S12: **Unsupervised nREM decoding: comparison with other decoders** (a) Average activity (average firing rate per neuron on y-axis) across the population (neuron index on x-axis, ordered according to their preferred angle from supervised tuning curve (TC)-based decoding of waking data), for a specific angle bin (Mouse 28; angle based on supervised decoding). Different panels correspond to different angle bins (15 bins in total). Blue, green, yellow: Waking, REM and nREM, respectively. (b) nREM points colored by SPUD, TC and population vector (PV) decoders. Insets on right: Top: difference in angles returned by SPUD and TC decoders. Middle: difference between angle returned by SPUD and TC decoders after removing times when population rate is $< 10$th percentile of waking activity. Bottom: difference between angle returned by SPUD and PV decoders after removing times when population rate is $< 10$th percentile of waking activity.

nREM manifold position over 200ms, and plot the mean LFP for the 5s before and after a large ($> 50$th percentile) or small ($< 50$th percentile) change, along with a 95% confidence interval computed as 1.96 times the standard deviation. For Fig. 4q, we convert the LFP to a spectrogram using a sliding Fourier transform, calculate the total power at each frequency in 1s windows, and correlate this with the summed absolute change in manifold position over 10 100ms bins (i.e. $\sum_{i=1}^{10} ||x(t_0+0.1*i)-x(t_0+0.1*(i-1))||$, where $t_0$ is the time at which the signals are being compared). We plot these correlations along with a 95% bootstrapped confidence interval, where we repeatedly resample the (LFP, change in manifold position) pairs and recompute the correlation.

To show that the lack of an observed ring manifold during nREM sleep is not due simply to lower firing rates during nREM when compared to waking, we subsample the waking spike counts to match the observed rates during nREM sleep. For neuron $i$, we define $p_i = \frac{\text{mean nREM rate}}{\text{mean waking rate}}$ and include each spike from neuron $i$ in our data independently with probability $p_i$. The ring manifold is clearly visible (Fig. S13a) in this subsampled data.

We apply SPUD to also decode from Mouse 25 during nREM sleep (decoding Mouse 12 during nREM is infeasible because of the confinement of the manifold to very low-activity states, Fig. S11). As with Mouse 28, we find that the SPUD angle agrees well with a tuning curve decoder, Fig. S13b,c. The dynamics when projected to the waking manifold are rapid and uncorrelated, rapidly settling to the equilibrium distribution, Fig. S13d,e. However, in the full population space, these dynamics show slower, correlated structure, Fig. S13f-h, including coherent rapid sweeps that are correlated with LFP power near 12 Hz, Fig. S13i.

### S13 nREM states and dynamics replicated in a continuous attractor model

To qualitatively reproduce states and dynamics during nREM, we add slow, large-amplitude fluctuations to the background and velocity inputs to the continuous attractor ring network model that was used to match waking and REM data.

As shown in Fig. S14a, the global amplitude fluctuations pull points off of the waking/REM manifold and toward the center, much as observed in the data. Also as in the data, the dynamics consist of a combination of staying in place and occasional large sweeps (Fig. S14a). In the model, slow global fluctuations in the background input (equivalently, the excitability) of neurons cause the population to intermittently return to the neighborhood of the zero activity state, thus resetting the angle dynamics every few hundred milliseconds and limiting the lifetime of the integrated representation. On the other hand, the large sweeps reflect large, slow fluctuations in velocity input when the network is not near the zero activity state. For instance, if the population that preferentially drives the activity bump clockwise has elevated activity for a few hundred milliseconds, the activity bump will sweep clockwise. The resulting network exhibits uncorrelated updates (Fig. S14b) and a diffusion plot with very high diffusivity (Fig. S14c) when projected to the waking manifold, but slower correlation dynamics in the full space, Fig. S14d. Moreover, epochs of large velocity changes are more coherent than epochs of small velocity changes, Fig. S14e, as in the real data. Thus, the previously described continuous attractor network with large, slowly fluctuating or tem-

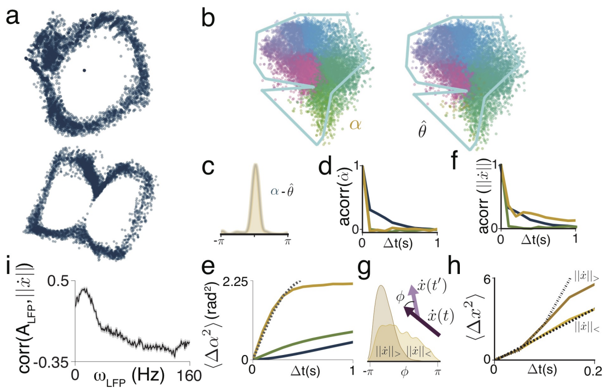


Figure S13: **Additional results on decoding, dynamics, and trajectory statistics during nREM.** (a) Testing the hypothesis that nREM manifolds appear higher-dimensional because of low spiking rates: Waking data only, but with spikes independently subsampled to match the mean firing rates observed during nREM. The low-dimensional ring manifold is still clearly visible. Panels (b-i) show results for Mouse 25. (b) nREM manifold colored by SPUD angle (left) and angle extracted from supervised decoding, based on waking tuning curves (right). (c) Histogram of differences in unsupervised (SPUD) and supervised angle estimates (the two estimates are first aligned by a global rotation and flip, as needed). (d) Autocorrelation of angular velocity (after projection of population state onto 1D waking manifold) during nREM (mustard); waking and REM velocity autocorrelations are in blue and green. (e) Plot of the average squared change in angle for different time separations. Dashed line: Diffusion on ring. Waking and REM shown for comparison. Note the much shorter autocorrelation time for projected nREM states. (f) Autocorrelation of velocity on full nREM manifold: note the appearance of much slower time-scales. Waking and REM angular velocity traces from panel (d) shown for comparison. (g) Histogram of angles between successive velocity vectors on full manifold (100ms separation): a peak at small angles indicates that the direction of motion along the manifold is correlated in time. Dark and light histograms for high and low velocity trajectories, respectively. Thus, high-velocity trajectories are more directionally coherent than low-velocity trajectories. (h) Diffusivity curves for states along full manifold, computed separately for high (dark) and low (light) velocity epochs, along with quadratic (coherent velocity-driven) and linear (random diffusion) fits. (i) Correlation of total change in position against LFP power in 1s bins.

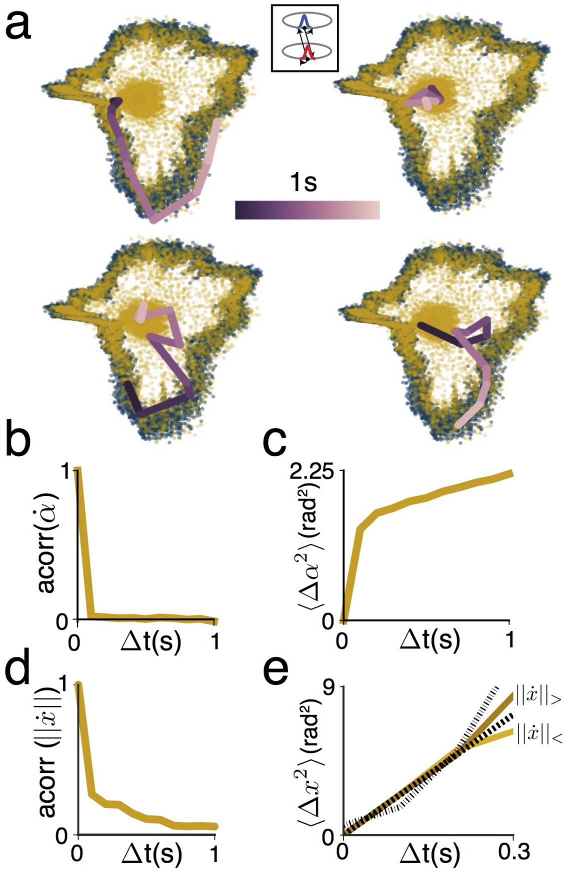


Figure S14: **Qualitative reproduction of nREM trajectory, dynamics, and statistics in continuous attractor ring network model.** (a) Sample trajectories in a model network with slow (temporally correlated) global fluctuations in the velocity and background inputs. nREM population states in mustard yellow; waking population states in blue for comparison (as in Fig. 4g). The states in the model show similar sweep-like and diffusing-in-place trajectories as the data. (b) Autocorrelation of the velocity of the population state after it is projected onto the 1D waking manifold (c) Diffusion plot for the projected population state shows a rapid increase in squared angle over time that quickly saturates. (d) Autocorrelation of velocity on full manifold (not projected onto 1D waking manifold). (e) Diffusion plot for consecutive large (dark) and small (light) velocity epochs on full manifold (defined as 300 ms of > 75th percentile or < 25th percentile of velocity distribution).

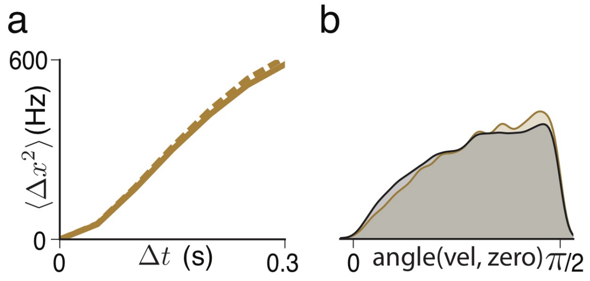


Figure S15: **nREM sweeps are not primarily driven by global rate fluctuations** (a) Sweeps from only high-rate epochs (dashed line) versus all sweeps (solid line). 84% of sweeps are preserved during high-rate epochs only, suggesting that sweeps are not primarily due to large global amplitude fluctuations. (b) Distribution of angles between the instantaneous velocity vector and the vector pointing toward the zero activity state (0 ($\pi/2$) means the instantaneous trajectory is toward (perpendicular to) the zero activity state), during sweeps (orange) versus at all times (black). Sweep trajectories are no more biased toward or away from the zero activity state than the dynamics at other times, again suggesting that sweeps are not preferentially center-in and center-out trajectories driven by global amplitude fluctuations.

porally correlated inputs qualitatively reproduces both the structure and the dynamics of the head-direction circuit during nREM sleep, and demonstrates how large global fluctuations can drive the representation off of the waking manifold and intermittently reset the encoded angle. The modeling results across states show how the dynamics for waking, REM and nREM can be reproduced using the same model and only changing the inputs.

The observation of global fluctuations in the nREM data suggests that the height of the attractor bump and thus the radius of the ring manifold in ADn is actually a neutral mode (rather than an intrinsic fixed point of the network dynamics), whose value is determined at least partially by external input. This is true in both excitation- and inhibition-dominated ring models, where neurons in the model network typically receive a fixed background input. It is will be interesting to understand how the equivalent background input or excitability is maintained at a fixed level across waking and REM (suggesting a form of fixed-point dynamics in the amplitude of the background input that persists in the transition from waking to REM), and how this set point is lost during nREM sleep.

For the nREM simulations, the velocity input is generated from a 0-mean Ornstein-Uhlenbeck process with correlation time 200 ms and standard deviation 1 rad/s. We truncate the velocity input at $\pm 2$ rad/s (i.e. values greater in magnitude than 2 are set to 2). We add background fluctuations to Eq. 3 that are of the form $\mu_0 + \mu_1 \cos(2\pi f_{bg} t)$, with $\mu_0 = -1.0$, $\mu_1 = 3.0$, $f_{bg} = 0.1$ Hz, and are clipped to lie below a maximum value of $-0.2$.

### S14 nREM sweeps are not primarily driven by global rate fluctuations

In Fig. 4m-o, we identify a regime of large coherent changes along the nREM manifold, which we call sweeps. To test whether these sweeps might simply reflect global fluctuations in population rate, we carry out two control analyses. First, we set a rate threshold at the bottom 20th percentile of the distribution of population rate activity (this retains 70% of nREM states), and only consider epochs where the nREM population firing rate stays above this threshold. We find that 84% of sweeps are preserved after this thresholding (thus sweeps are overrepresented during high rates), and that the diffusion curve for this subset of sweeps is near-identical to that of all sweeps, Fig. S15a. Second, at each moment in time we calculate the angle between the instantaneous velocity vector and the vector toward the zero-activity state. If sweeps preferentially occur during global rate fluctuations, then these vectors should be aligned. We find that velocities during sweeps are not preferentially directed towards or away from the zero activity state, Fig. S15b.

### References


[1] Peyrache, A., Lacroix, M. M., Petersen, P. C. & Buzsáki, G. Internally organized mechanisms of the head direction sense. *Nat. Neurosci.* **18**, 569–575 (2015).

[2] Bartlett, M. The square root transformation in analysis of variance. *Supplement to the Journal of the Royal Statistical Society* **3**, 68–78 (1936).

[3] Munkres, J. R. *Topology* (Prentice Hall, Upper Saddle River NJ, 2000).

[4] Tenenbaum, J. B., De Silva, V. & Langford, J. C. A global geometric framework for nonlinear dimensionality reduction. *Science* **290**, 2319–2323 (2000).

[5] Kingma, D. P. & Welling, M. Auto-encoding variational bayes. *Proceedings of the International Conference on Learning Representations (ICLR)* (2014).

[6] Rezende, D. J., Mohamed, S. & Wierstra, D. Stochastic backpropagation and approximate inference in deep generative models. *Proceedings of the 31st International Conference on Machine Learning, pp. 12781286* (2014).

[7] Roweis, S. T. & Saul, L. K. Nonlinear dimensionality reduction by locally linear embedding. *Science* **290**, 2323–2326 (2000).

[8] Ghrist, R. Barcodes: The persistent topology of data. *Bull. Amer. Math. Soc.* **45**, 61–75 (2008).

[9] Carlsson, G. Topology and data. *Bull. Amer. Math. Soc.* **46**, 255–308 (2009).

[10] Singh, G. *et al.* Topological analysis of population activity in visual cortex. *J Vis.* **8**, 1–18 (2008).

[11] Curto, C. & Itskov, V. Cell groups reveal structure of stimulus space. *PLoS Comput. Biol.* **4**, e1000205 (2008).

[12] Bubenik, P. Statistical topological data analysis using persistence landscapes. *J. Mach. Learn. Res.* **16**, 77–102 (2015).

[13] Kwitt, R., Huber, S., Niethammer, M., Lin, W. & Bauer, U. Statistical topological data analysis-a kernel perspective. In *Advances in Neural Information Processing Systems 28 (NIPS)*, 3070–3078 (2015).

[14] Softky, W. R. & Koch, C. The highly irregular firing of cortical cells is inconsistent with temporal integration of random EPSPs. *J. Neurosci.* **13**, 334–350 (1993).

[15] Shadlen, M. N. & Newsome, W. T. The variable discharge of cortical neurons: implications for connectivity, computation, and information coding. *J. Neurosci.* **18**, 3870–3896 (1998).

[16] Kuffler, S. W., Fitzhugh, R. & Barlow, H. B. Maintained activity in the cat's retina in light and darkness. *J. Gen. Physiol.* **40**, 683–702 (1957).

[17] Goris, R. L., Movshon, J. A. & Simoncelli, E. P. Partitioning neuronal variability. *Nat. Neurosci.* **17**, 858–865 (2014).

[18] Bauer, U., Tralie, C. & Saul, N. Ripser. *Github* (2017). URL `https://github.com/ctralie/ripser`.

[19] Grassberger, P. & Procaccia, I. Measuring the strangeness of strange attractors. *Physica D* **9**, 189–208 (1983).

[20] Hafting, T., Fyhn, M., Molden, S., Moser, M.-B. & Moser, E. I. Microstructure of a spatial map in the entorhinal cortex. *Nature* **436**, 801–806 (2005).

[21] Yoon, K. *et al.* Specific evidence of low-dimensional continuous attractor dynamics in grid cells. *Nat. Neurosci.* **16**, 1077–84 (2013).

[22] Burak, Y. & Fiete, I. R. Fundamental limits on persistent activity in networks of noisy neurons. *Proc. Natl. Acad. Sci. U.S.A.* **109**, 17645–50 (2012).
